# Partitioned Local Depth analysis of time series transcript abundance data reveals elaborate community structure

**DOI:** 10.1101/2022.07.25.501352

**Authors:** Maleana G. Khoury, Kenneth S. Berenhaut, Katherine E. Moore, Edward E. Allen, Alexandria F. Harkey, Joëlle K. Muhlemann, Courtney N. Craven, Jiayi Xu, Suchi S. Jain, David J. John, James L. Norris, Gloria K. Muday

## Abstract

Transcriptome studies that provide temporal information are valuable for identifying groups of similarly-behaving transcripts, giving insight into overarching gene regulatory networks. Nevertheless, inferring transcriptional networks from time series data is challenging, in part because it is difficult to holistically consider both local relationships and global structure of these complex and overlapping transcriptional responses. To address this need, we employed the Partitioned Local Depth (PaLD) method to examine four time series transcriptomic datasets generated using the model plant *Arabidopsis thaliana*. Here, we provide a self-contained description of the method and demonstrate how it can be used to make predictions about gene regulatory networks based on time series data. The analysis provides a global network representation of the data from which graph partitioning methods and neighborhood analysis can reveal smaller, more well-defined groups of like-responding transcripts. These groups of transcripts that change in response to hormone treatment (e.g., auxin or ethylene) or high salinity were demonstrated to be enriched in common biological function and/or binding of transcription factors that were not identified with prior analyses of this data using other clustering and inference methodologies. These results reveal the ability of PaLD to generate predictions about gene regulatory networks using time series transcriptomic data, which can be of value to the systems biology community.

## Introduction

Transcriptome remodeling is a highly dynamic, tightly controlled process during which transcript abundance is altered in response to stimuli (e.g., hormones and stress responses) [1–10] with precise temporal kinetics. Insight about the transcriptional regulatory networks downstream of stimuli can be extracted from time series data, which provides information about changes in transcript abundance over time [11–16]. Determining not only which genes have altered transcript abundance in response to stimuli but also inferring which of these genes might operate in the same transcriptional networks based on common temporal patterns provides greater depth to our understanding of signaling pathways driven by transcriptional cascades. Genes that have the same temporal dynamics form subnetworks that may be turned on by the same upstream transcription factors (TFs) or turned off by the same repression mechanisms. Additionally, common temporal kinetics for transcript synthesis may allow coordinated responses if subnetworks include transcripts encoding proteins that function in the same biological processes [17–22]. Time series data can also be used to infer the sequential activation of genes in a network using next-state modeling, which combines information about TF-target interactions with information about the temporal dynamics of gene expression [23–27]; a rapid increase in the synthesis of TFs suggests that these proteins may control the expression of downstream target genes [28–30]. Overall, transcriptome studies that measure changes in transcript abundance over time provide valuable information about genes that are co-expressed or are sequentially activated, information that can be used to predict gene regulatory networks [31–34].

Inferring transcriptional networks from time series transcriptomic data remains a challenge for biologists. Commonly employed clustering and labelling methods, such as hierarchical clustering and *k*-means clustering [35,36], often require additional user input (e.g., defining the number of clusters) that might not fit the naturally occurring pattern of clusters in the data. In addition to these tools, other methods have been developed to infer gene networks from time series data [37–45]. At their core, these techniques group entities (or objects) together into classes (or collections of objects). While clustering techniques often provide class labels at a local level, they do not provide information on the precise connections between the underlying entities (transcripts) and between classes (clusters). Network inference methods that can reveal information not only about local connections but also global relationships with minimal user input are needed to develop better models for predicting gene networks. Recently, Berenhaut, Moore, and Melvin introduced a new technique, Partitioned Local Depth (PaLD), that provides both local- and global-level information when examining social science datasets [46]. This study is the first application of PaLD to transcriptomic data and this approach to reveal connections between genes in the form of a weighted network, in which edge weights (and resulting proximity in the network) convey relationship strengths between data points. This approach is motivated by a community-level social perspective in which an underlying assumption is that an object (transcript) is relatively similar to its *nearby* neighbors. Here, the concept of “nearby” reflects local perspective and adapts to varying density without the need for extraneous parameters that restrict the number of groups, such as the optimization that is required to determine the number of K-means clusters [47]. Community detection methods can then be applied to the weighted networks obtained from PaLD to reveal smaller groups of transcripts with the most similar responses, but that maintain connectivity to the overall structure, thereby providing a holistic view of the patterns and connections within and between community groups [48,49].

In this study, we applied PaLD to search for relationships between transcripts whose abundance changes in several time series microarray or RNA-Seq datasets from the model organism *Arabidopsis thaliana*. We then evaluated the efficacy of PaLD in this context by asking whether groups of like-responding transcripts had conserved function using Gene Ontology (GO) annotations, and whether these groups had conserved regulation by uncovering TFs binding to promoter regions of genes in these groups. The publicly available datasets chosen for this analysis measured changes in transcript abundance across a broad time series. Two of these datasets resulted from treatments that elevated the levels of the gaseous plant hormone ethylene, with transcript changes defined by oligonucleotide microarray analyses [50] or RNA-Seq [51]. Two other datasets examined transcriptional responses to elevated levels of auxin [52] or salt [53]. These datasets were chosen because of the extensive sampling across time and because both auxin and salt have been shown to interact with ethylene to modulate root development [54,55]. We showed that PaLD can reveal patterns in time series data by grouping transcripts with closely related temporal responses, revealing subnetworks of transcripts that were also enriched in common biological processes and are targets of the same TFs. Furthermore, the combinations of functional enrichments and TF targets were subnetwork*-*specific, meaning these unique combinations were not shared between groups. Although the intention of PaLD is to provide additional insights beyond just sorting genes into like-responding groups, we compared the PaLD groups to clusters and groups identified by other approaches, including *k-*means clustering and the Short Time Series Expression Miner (STEM) [50,52,53], and showed that PaLD can reveal new groups of transcripts that are enriched in biological processes and targets of TFs not previously identified in the data. These findings demonstrate the utility of PaLD as a transparent and effective method to search for patterns within time series transcriptomic data and illustrate how it can be used to generate novel predictions about gene regulatory networks.

## Algorithm

### Description of the PaLD Method

The PaLD method looks for community structure using dissimilarity comparisons. Suppose that *S* is a set of *n* points (i.e., |*S*| = *n*) and write *x* ∈ *S* to denote membership in *S*. For each pair of points, *x*, *y* ∈ *S*, we denote the dissimilarity (or distance) from *x* to *y* by *d*(*x*, *y*). Throughout, the only information leveraged is dissimilarity comparisons of the form *d*(*z*, *x*) < *d*(*z*, *y*) or *d*(*x*, *z*) < *d*(*y*, *z*), for *x*, *y*, *z* ∈ *S*. The foundation in distance comparisons allows for an element of robustness and a broad interpretation of distance; the only metric requirement is that each *x* be closer to itself than to any other point. For example, one may employ application-specific measures of dissimilarity based on correlation coefficients or fixation indices, in which dissimilarity is, say, interpreted non-linearly. The results are unchanged by any re-scaling of the distances (e.g., logarithmic or exponential).

Embedding *S* in a latent (social) space in which local comparisons (or “conflicts”) arise, we define a measure of pairwise relationship strength by considering alignment in induced local regions (or foci). This perspective allows for consideration of interactions occurring across a variety of scales and gives a meaning of *local* that does not require the user to provide additional inputs or search an underlying parameter space. We present the algorithm in a concrete computational form, but in the note below, we briefly describe the probabilistic social perspective from which the PaLD approach arises; for further details see [46].

For fixed *x*, *y* ∈ *S*, define the *local focus*, *U*_*x*,*y*_, to be the set of points which are as close to *x* as *y* is to *x*, or as close to *y* as *x* is to *y*, that is,

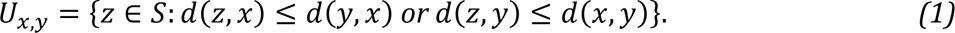

We consider a point *w* ∈ *S* to be aligned with *x* in opposition to *y* if *w* is in the local focus (i.e., *w* ∈ *U*_*x*,*y*_) and is closer to *x* than to *y* (i.e., *d*(*w*, *x*) < *d*(*w*, *y*)). The strength of the alignment is then inversely proportional to the number of points in the local set (i.e., |*U*_*x*,*y*_|). The *cohesion* of *w* to *x* is defined via

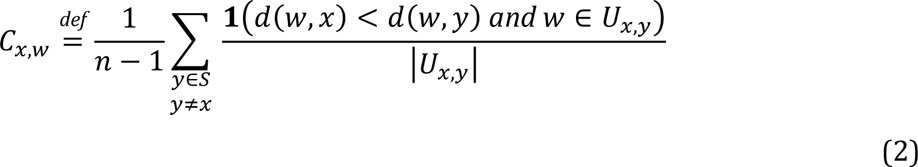

i.e., it is the average strength of alignment, where |*U*_*x*,*y*_| is the number of elements in the set *U*_*x*,*y*_ and **1**(*Q*) = 1 if statement *Q* is true and **1**(*Q*) = 0, otherwise. In the case that *d*(*w*, *x*) = *d*(*w*, *y*), we resolve the tie by setting the numerator to 1/2; for clarity of presentation this case is suppressed.

To identify important relationships between points, we will emphasize those pairs for which mutual cohesion is greater than what is typical for a point in a local focus. Leveraging the symmetry of the local foci, we consider the relationship between *x* and *w* to be (*particularly) strong* whenever

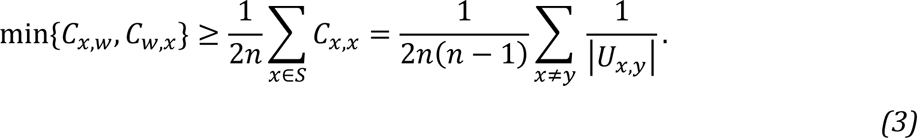

Further context and justification for the value of this threshold from a probabilistic perspective are given in (Berenhaut et al. 2022).

For *S* = {*s*_1_, *s*_2_, …, *s*_*n*_}, we obtain an *n* × *n* matrix of cohesion values, ***C*** = [*C*_*si*,*sj*_]. The cohesion matrix conveys strength of relationship between points and can be visualized as a network (or graph). The adjacency matrix defining the weighted undirected graph, *G*, is obtained by replacing each value, *C*_*si*,s*j*_, with the minimum of *C*_*si*,s*j*_ and *C*_*sj*,s*i*_. Thresholding the symmetrized cohesion matrix at the value in Eq. (3), leads to an adjacency matrix defining the graph, *G*^*^, whose (weighted) edges are only those for which the cohesion is particularly strong.

For visualization purposes, we apply the Kamada-Kawai graph drawing algorithm [56] to the graph *G*^*^. When further partitioning is desired (for instance, as in Section 3.2), we may employ a community detection algorithm, such as the Louvain algorithm, on the largest connected component. Louvain is one of several algorithms available to find a partition of a network into groups which maximizes modularity.

***Note:*** (Probabilistic perspective and local depth). Aggregating the support for *x* from all points *w* ∈ *S*, we obtain a measure of local centrality, referred to as *local (community) depth*, and defined by

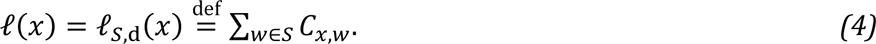

From the probabilistic perspective considered in [46], the local depth of *x*, ℓ(*x*), described above is the probability that, in opposition with a point *Y* selected uniformly at random from the remaining points in *S*, a focus-point, *Z* ∈ *U*_*x*,*Y*_ selected uniformly at random is closer to *x* than it is to *Y*. We then have the following,

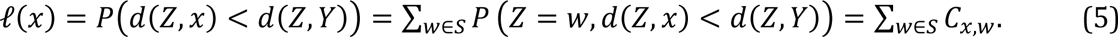

In particular, cohesion, as equivalently defined in Eq. (2), is obtained from partitioning the probability which defines local depth; thus, this approach is entitled *Partitioned Local Depth*.

**Example** (Example 1, revisited). Consider the distances in Fig. 1A. To compute the cohesion of *s*_4_ to *s*_2_, we consider the local foci for *j* = 1,3,4,5,6,7. Among these 6 induced sets, *s*_4_ is contained in *U*_*s*2,*s*1_ = *S*, *U*_*s*2,*s*4_ = *U*_*s*2,*s*5_ = {*s*_2_, *s*_3_, *s*_4_, *s*_5_}, *U*_*s*2,*s*6_ = {*s*_1_, *s*_2_, *s*_3_, *s*_4_, *s*_5_, *s*_6_} and *U*_*s*2,*s*7_ = *S*. Across these sets, *s*_4_ is closer to *s*_2_ than to the opposing point, *s*_*j*_, for *j* = 1,6,7. As a result, as in Eq. (2),

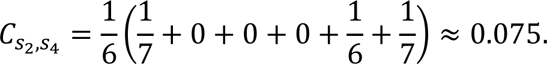

**Fig. 1.**
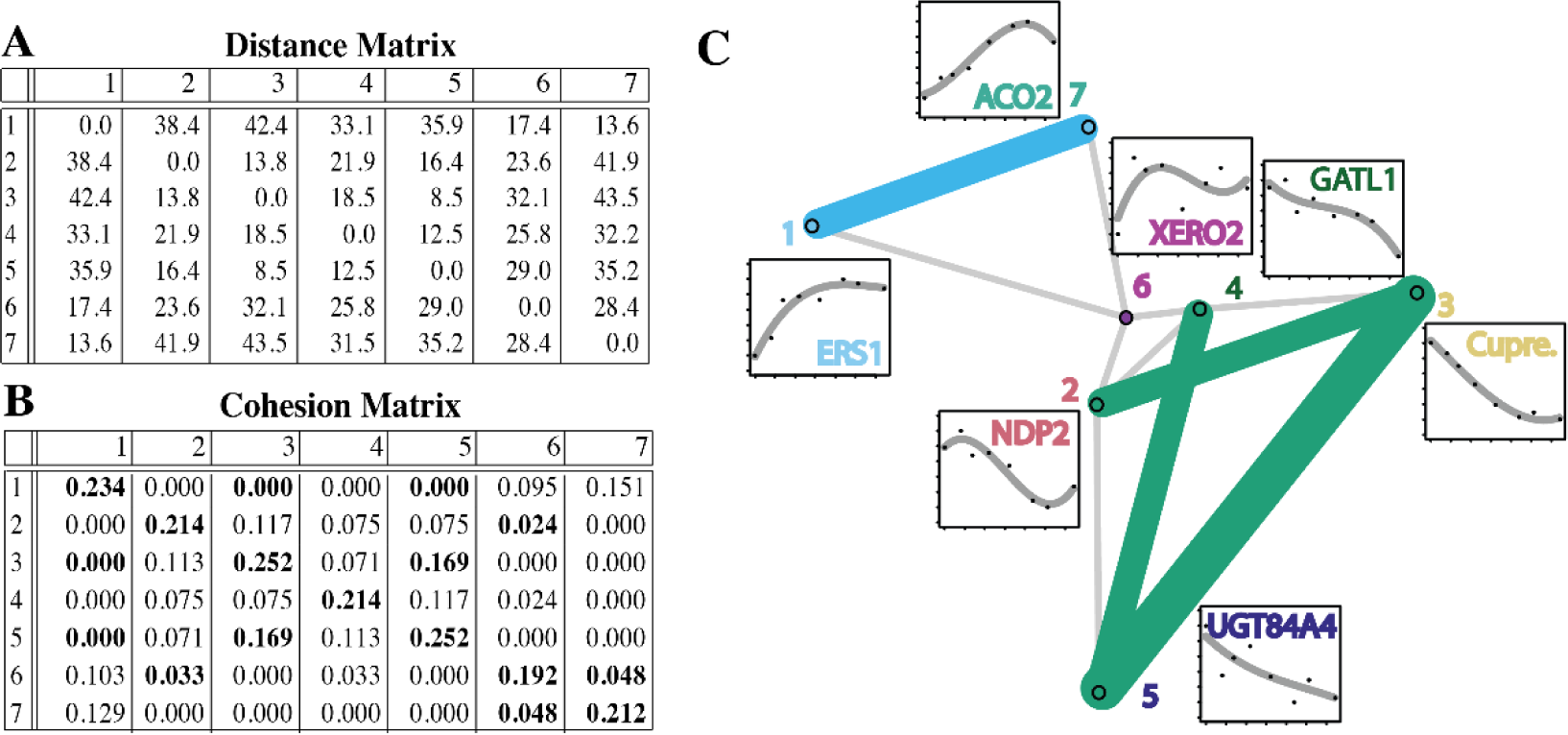
An illustrative example of PaLD applied to a small set of 7 transcripts examined across 8 time points. In A, the matrix of distances between transcripts is given. In B, the matrix of cohesion values is displayed, see *Eq. (2)*]. Larger values of cohesion indicate stronger pairwise relationships; those which are particularly strong (*Eq. (3)*) are indicated in bold. In C, the resulting community graph is displayed together with insets that show the temporal response of the transcripts. Dots represent the average min-max scaled signal log ratios across three biological replicates at each time point used in the study.

The remaining pairwise cohesion values are computed similarly and are displayed in Fig. 1B. The threshold distinguishing particularly strong relationships, here, is 0.112 (i.e., half the average of the entries on the diagonal; see Eq. (3)). Relationships which are particularly strong are indicated in bold in the cohesion matrix in Fig. 1B. In Fig. 1C, we display the graph *G*; the subgraph graph, *G*^*^, of strong relationships is emphasized with bold colored edges.

### Defining a Distance between Transcripts

A goal of this study is to reveal network-based (local and global) relationship information among transcripts, from the shape of the curves associated with expression level over time. To apply the method described above, we require an informative sense of dissimilarity between transcripts. Here, we take the average at each time point over trials, and then apply min-max scaling over the relevant time points. A min-max scaling linearly transforms the data to a [0, 1] range. Each of the datasets considered here were collected at less than 10 approximately exponentially spaced time points; to reflect decay in activity we consider the data in log time. Using least squares, we construct cubic polynomials, *f*_*i*_(*t*), 1 ≤ *i* ≤ *n* (one for each gene), to model the double-extrema behavior of interest. To capture other behaviors, different non-linear or piecewise-linear functions could be used. To measure the distance between the fitted time-curves, *f*_*i*_ and *f*_*j*_, (associated to two different genes), we consider similarity in functional value and first derivative (capturing the expression level and rate of change, respectively). In particular, we define the associated distance between the 2 curves via

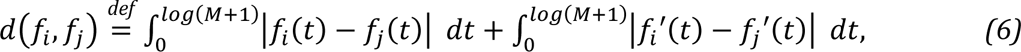

where 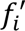 is the first derivative of *f*_*i*_ and *M* is the maximal time point. Note that the first term is the total (absolute) area between the curves; and the second corresponds to the (absolute) area between the derivatives. Throughout, the distance between transcript abundance profiles is understood to be the dissimilarity of the associated fitted cubic polynomials as described in Eq. (6). The distance function given in Equation (6), defines the distance between 2 curves as the area between the graphs of the 2 curves plus the area between the graphs of their derivatives. There are many possible distance metrics, depending on the application at hand. For instance, standardizing the coefficient of 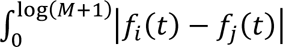 as 1, yields distance functions

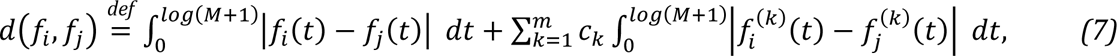

where *m* is a positive integer, {*c*_*i*_} are real coefficients and *f*^(*k*)^ is the *k*^*t*ℎ^ derivative of *f*, are all distance functions. By changing the weights, {*c*_*k*_}, one can choose to emphasize or de-emphasize the value of fundamental properties of the corresponding graph. In our example, we chose to include the area between the graphs of the derivatives so as to give additional weight reflecting whether the corresponding functions were increasing or decreasing similarly. By leaving out the term corresponding to the second derivative, we chose to ignore the effect of concavity. The number of time points in the dataset as well as the predicted underlying curves are important considerations in determining the definition of the corresponding distance function. The definition of appropriate distance metrics for the data is an important open question.

### An Illustrative Example

In Fig. 1, for illustration, we apply the PaLD method to reveal community structure in a subset of 7 transcripts from a larger microarray dataset [50]. The sole input for PaLD is a set of pairwise distances (or, more generally, distance comparisons). The pairwise distances between the transcripts are displayed in Fig. 1A.

The output cohesion matrix, indicating relationship strength, is provided in Fig 1B, and the resulting community structure is displayed in Fig 1C. Relationships which are particularly strong (see Eq. (3) <u>below)</u> are indicated in bold in Fig. 1B and emphasized with wider colored edges in Fig. 1C. For more details on these computational methods, see [46].

### Enrichment Analysis Pipeline

Four publically available time-series transcriptomic datasets were used in our analysis. For more information about the treatment conditions and time points used in each of those studies, see Supplemental Methods and Table S1.

Identification of enriched binding of transcription factors in the clusters and groups was achieved using TF DEACoN, a program that uses DAP-Seq data [57] to report targets of a subset of transcription factors in Arabidopsis [58]. The size of some groups is larger in the enrichment analysis due to microarray probes recognizing transcripts from more than one gene or smaller due to probes that were not associated with a gene identifier being removed. The output of this analysis uses a log fold change cutoff greater than 1 and a *p*-value cutoff of less than 0.05 and is such that the abundance in the group is compared to the whole genome. (Note that adjusted *p*-values were used here, yielding slightly different results than those obtained using raw values.)

GO enrichment in PaLD groups, *k-*means clusters, or STEM groups, relative to the whole genome, was performed in the Singular Enrichment Analysis tool in AgriGO [59]. In AgriGo, “statistical test method” was set to hypergeometric and “multi-test adjustment method” was set to Yeketuli (FDR under dependence). The top 3 most significantly enriched GO annotations were reported. All GO terms reported had adjusted *p*-values less than or equal to 0.05. We also used Metascape [60] to identify a list of GO terms associated with each gene in the IAA microarray dataset. Metascape used a comprehensive list of knowledgebases, and therefore was more appropriate for our search [60]. We analyzed against the Arabidopsi*s* genome and used the Express Analysis tool [60].

### Implementation

#### PaLD analysis of an ACC microarray dataset yielded a network with local and global connections

The PaLD method was used to generate networks using four different datasets. The details on these datasets are provided in the supplemental materials. These included two microarray datasets, which identified genome wide changes in roots of *Arabidopsis thaliana* after treatment with the ethylene precursor ACC [50], the auxin indole-3-acetic acid [52], and two RNA-Seq datasets in Arabidopsis seedlings treated with ethylene gas [51] or elevated levels of salt [53].

We began by applying PaLD to generate a network using 447 transcripts whose abundance changed in response to treatment with 1-aminocyclopropane-1-carboxylic acid (ACC), which is the immediate precursor to the plant hormone ethylene. This treatment increases ethylene synthesis and signaling [50]. The resulting network structure illustrated local and global relationships among the transcripts, which is shown with each node representing a gene and each edge connecting the transcripts with a locally similar temporal response (Fig. 2). The network contained 2 large groups of transcripts that were largely downregulated (at the top of the diagram) or upregulated (in the lower part of the diagram), while a smaller group in the middle of the network contained transcripts with mixed responses. We selected 7 transcripts (the same transcripts used in the illustrative example) to represent different parts of the network. For each transcript, we generated a fiber plot, which contains fitted curves reporting the abundance of those 7 transcripts as a function of time after ACC treatment (Fig. 2). Although visually similar, these fiber plots are not to be confused with Andrews plots for displaying multivariate structure (see [61]).

**Fig. 2.**
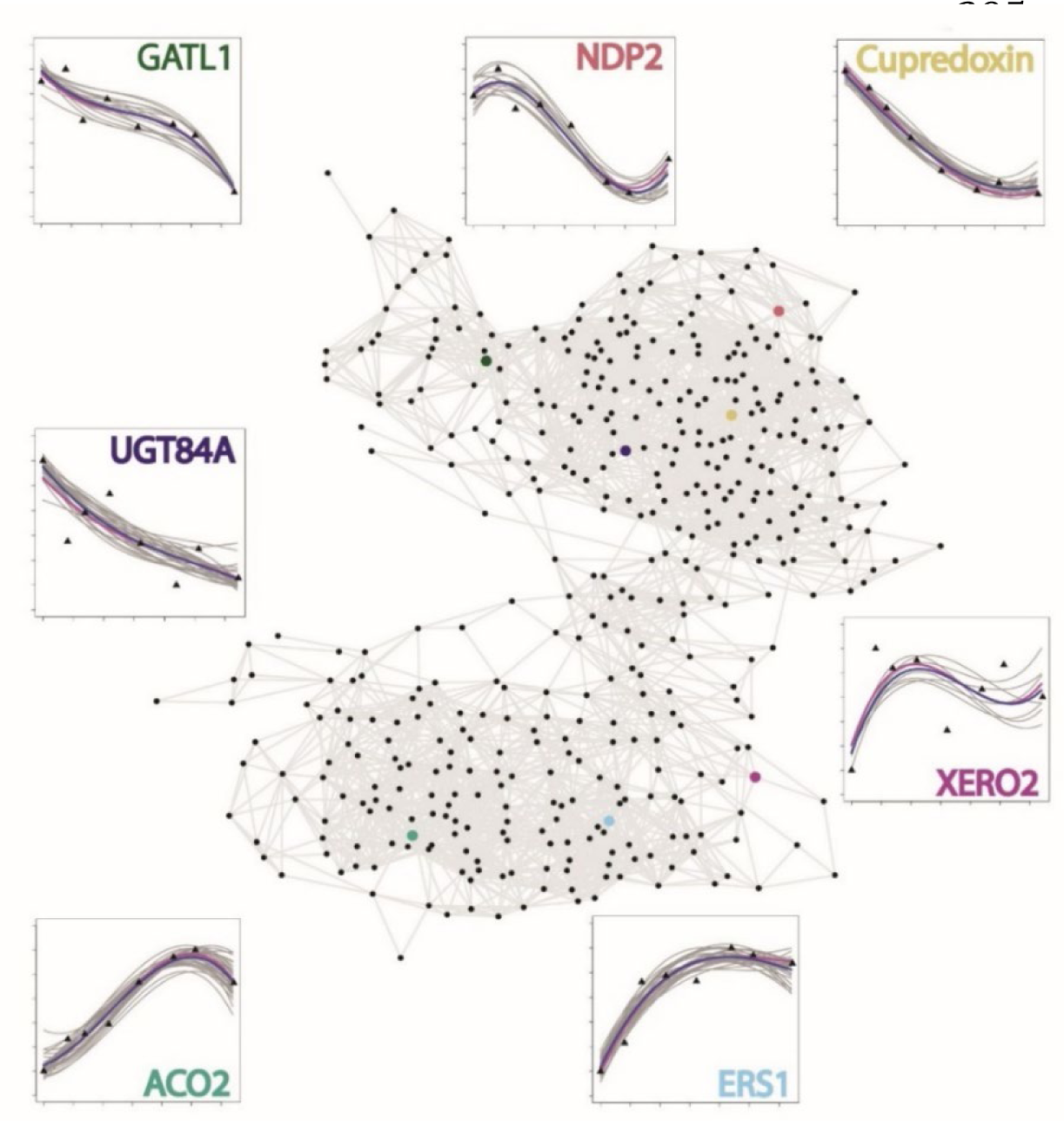
PaLD network of the 447 transcripts in the ACC microarray dataset [50]. Each node represents a single transcript, and each edge represents a pairing between nearest neighbors. Fiber plots illustrate transcript abundance over time for 7 representative genes (magenta lines) and their nearest neighbors (gray lines) averaged across 3 biological replicates, with the position of each transcript in the network indicated in the same color as the gene names. Averages for each neighborhood are shown as blue lines. Black triangles indicate the average abundance at each time point. Data is considered in log time and has been min-max scaled.

Additionally, smaller subnetworks, or neighborhoods, were determined for each of these 7 transcripts. These neighborhoods consisted of a transcript and all of its direct connections in the network, which are shown in gray on the fiber plots. These results illustrated the strength of PaLD in grouping genes with similar behavior, as the fibers in each of these plots followed the same trends. The varying neighborhood sizes reflecting local structure is a key feature of PaLD not exhibited by naïve nearest neighbor methods (see [46]).

To divide this network into larger neighborhoods of like-responding genes, we used the Louvain method for community detection [15]. This method maximizes modularity, a quantity that is larger for partitions with stronger within-group ties relative to that which would be expected by chance. When used in combination with PaLD, this approach revealed 7 regions of temporally similar transcripts, which are shown with color coded nodes on a network diagram (Fig. 3A). The transcripts in each of these PaLD groups are listed in Supplemental Dataset 1, along with the abundance changes in these transcripts reported as the signal log ratio (SLR) over the time course of ACC treatment. Fiber plots of transcript abundance over time, with the averages shown in blue and the behavior of the corresponding genes shown in gray, were overlaid on the network revealing that this method identifies patterns of transcripts with similar kinetic response (Fig. 3B). The transformed gene expression data prior to fitting the cubic polynomials illustrated that group 6, which contained transcripts with mixed responses, had the highest variability in temporal gene expression pattern of all the PaLD groups (Fig. S1).

**Fig. 3.**
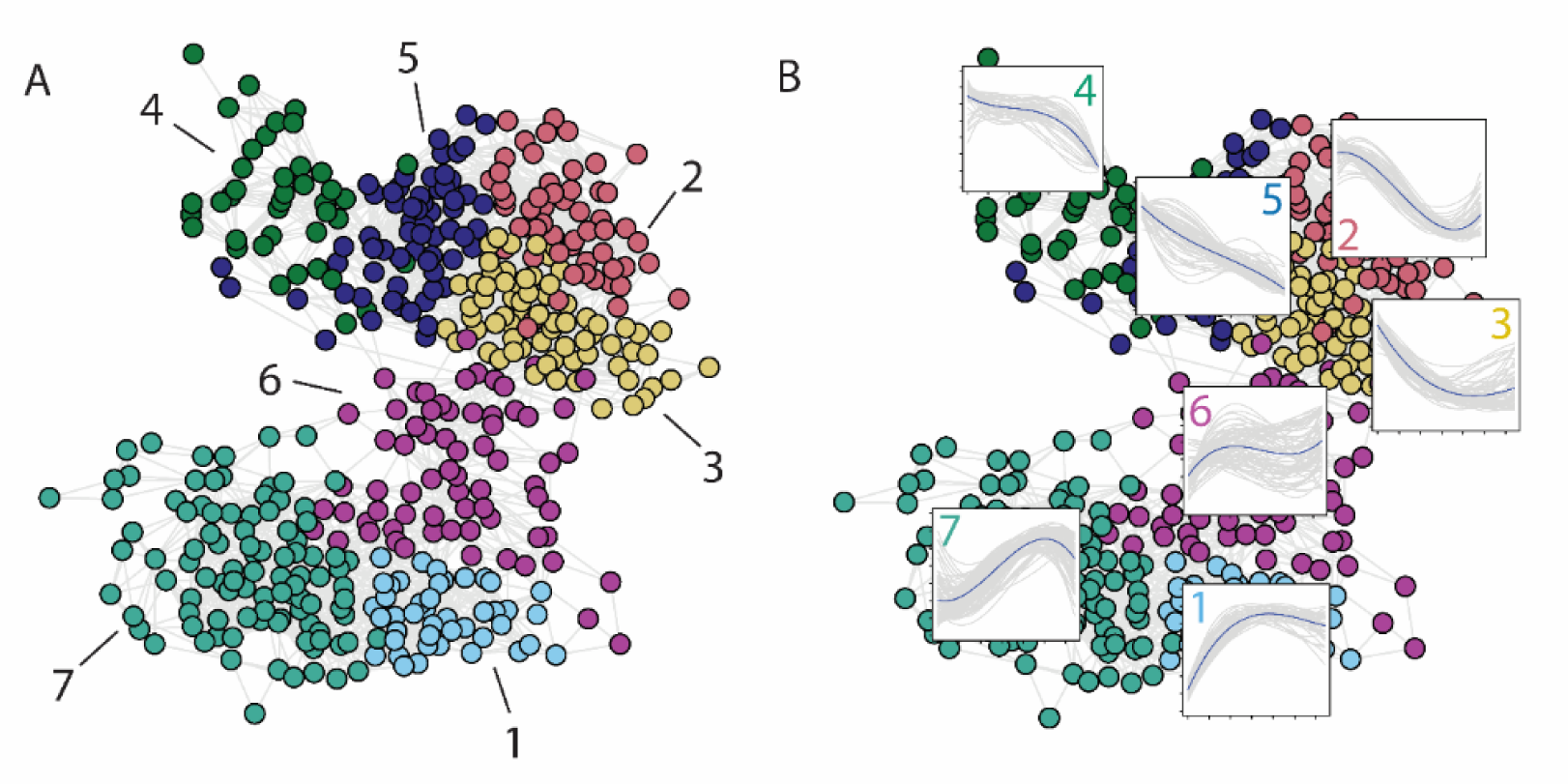
PaLD was used in combination with a community detection algorithm to partition the ACC microarray dataset into larger groups with similar kinetic responses. (A) Seven groups, or subnetworks, were revealed by application of the Louvain method for community detection to the PaLD network. Each group is color coded and labeled 1–7. (B) Fiber plots illustrate temporal responses of all transcripts within a group with the averages of all transcripts in that group shown in blue.

#### Comparison of subnetworks predicted by PaLD to those predicted by *k*-means clustering

One goal of this work is to understand how PaLD groups containing transcripts with similar kinetic response compare to methods used to sort transcripts into like-responding groups or clusters. To evaluate the ability of PaLD to organize transcripts as compared to the frequently used *k*-means clustering, we compared the PaLD groups to previously generated *k-*means clusters using this same set of data. The *k-*means clustering method was previously applied to this group of 447 transcripts yielding 72 clusters ranging in size from 1 to 111 genes, with 48 clusters of size 1 and 24 clusters of size greater than or equal to 2 [50]. The transformed gene expression data for transcripts in each of the top ten *k*-means clusters prior to fitting the cubic polynomials are shown in Fig. S2. To visually compare the *k-*means clusters to PaLD groups, we colored the nodes in the PaLD network to correspond to each of the 10 largest *k-*means clusters, which we refer to as A-J (corresponding to Clusters 1-10 in [50]), with this network shown in Fig. 4.

**Fig. 4.**
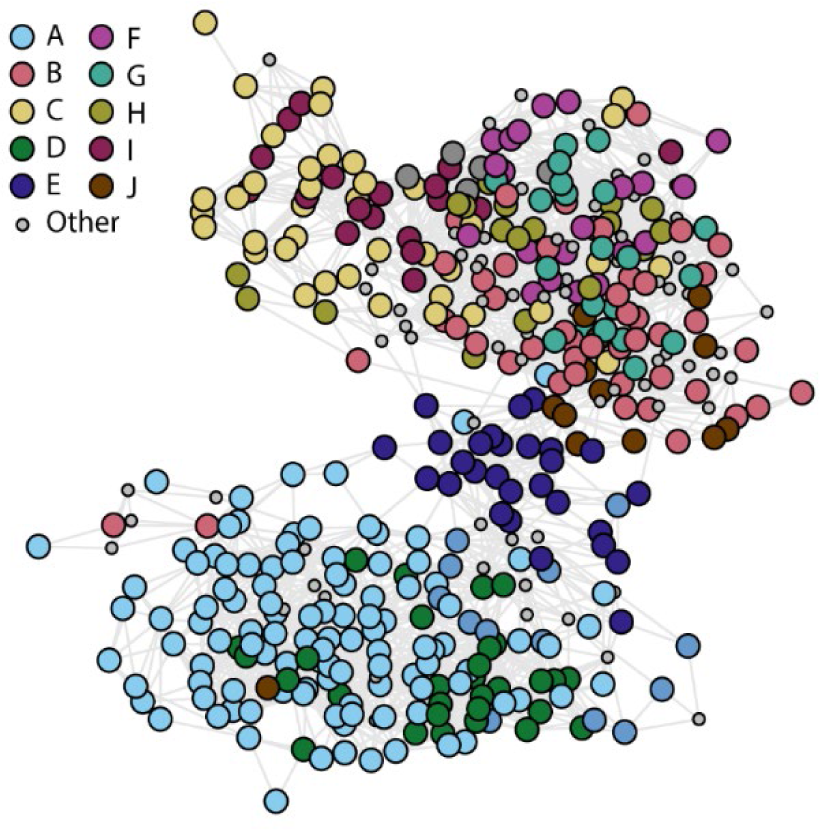
Clusters of like-responding transcripts identified by k-means clustering were overlaid on the PaLD ACC network. The position of transcripts from the top 10 k-means clusters determined previously [50] were denoted by distinct colors within the PaLD ACC network. Transcripts that were not part of the top 10 k-means clusters are shown as smaller gray circles in the network.

We also quantitatively compared the *k-*means clusters to the groups previously defined for the PaLD network using a table to illustrate overlaps between PaLD groups and *k-*means clusters (Table 1). The *k-*means clusters A, D, and E had the least spread across the PaLD groups, which is consistent with the diagram in Fig. 4. The matrix illustrates that 100% of the genes in cluster E were in PaLD group 6, 72% of the genes in cluster A (80/111 transcripts) were in group 7, and 67% of the genes in cluster D were in group 1, while other PaLD groups showed broader distribution between *k-*means clusters, with cluster J containing genes in 5 PaLD groups with no more than 33% of those genes in any single PaLD group.

**Table 1.** A comparison of PaLD groups and the k-means clusters in [50]. The PaLD approach revealed 7 groups while 18 k-means clusters were originally considered (the largest 10 highlighted here). PaLD groups and k-means clusters (of size less than 15) are combined as “Other” in the table (comprising 60 k-means clusters with a total of 105 transcripts and 7 PaLD groups with a total of 8 transcripts, respectively). As an example, PaLD group 3 consists of 1 element of k-means cluster A, 24 elements of B, 2 elements of C, etc. The “Other” category is much larger for k-means as many transcripts were not in the top 10 clusters and had fewer than 15 transcripts.

|  |  | *k*-means Clusters |  |  |  |  |  |  |  |  |  |  |  |
| --- | --- | --- | --- | --- | --- | --- | --- | --- | --- | --- | --- | --- | --- |
|  |  | A | B | C | D | E | F | G | H | I | J | Other | Sum |
| PaLD Groups | 1 | 16 | 0 | 0 | 20 | 0 | 0 | 0 | 0 | 0 | 0 | 7 | 43 |
|  | 2 | 0 | 10 | 4 | 0 | 0 | 9 | 9 | 5 | 1 | 2 | 16 | 56 |
|  | 3 | 1 | 24 | 2 | 0 | 0 | 5 | 10 | 2 | 0 | 5 | 20 | 69 |
|  | 4 | 0 | 0 | 25 | 0 | 0 | 0 | 0 | 0 | 10 | 0 | 15 | 40 |
|  | 5 | 0 | 10 | 8 | 0 | 0 | 8 | 2 | 11 | 7 | 0 | 17 | 63 |
|  | 6 | 13 | 0 | 0 | 4 | 28 | 0 | 0 | 0 | 0 | 5 | 19 | 69 |
|  | 7 | 80 | 2 | 0 | 6 | 0 | 0 | 0 | 0 | 0 | 1 | 10 | 99 |
|  | Other | 1 | 2 | 0 | 0 | 0 | 0 | 0 | 0 | 0 | 2 | 3 | 8 |
| Sum |  | 111 | 48 | 39 | 30 | 28 | 22 | 21 | 18 | 18 | 15 | 105 | 447 |

We also compared the unfitted curves of PaLD groups 1 and 7 to clusters D and A, respectively, to illustrate the more temporally linked gene expression patterns of the transcripts within the PaLD groups (Fig. S3). This analysis revealed that some sets of transcripts have consistent behavior and are grouped together by both *k-*means clustering and PaLD, while for other transcripts these methods detect relationships in different ways.

#### Enrichment analysis of PaLD groups in the ACC microarray dataset revealed conserved biological functions and transcriptional regulators

We hypothesized that transcripts which respond similarly in time may work together to drive the same biological processes and that their synthesis may be regulated by the same TFs. Since genes often work together in smaller subnetworks, we reasoned that the groups revealed by the community detection algorithm would be large enough to yield enrichment results yet small enough to represent biologically meaningful subnetworks. Therefore, to assess the ability of PaLD to predict genes that function, and are regulated, in the same biological network, we examined whether there were enriched functions and TF targets within the PaLD groups (Table 2). Using AgriGO v2.0, we identified enriched function in these groups, finding that 6 out of 7 PaLD groups were significantly enriched in GO annotations. To examine enriched targets of TFs in each of the groups, we used TF DEACoN [58], which identifies significant enrichment in genes that have been shown to be targets of TFs defined by a publicly available DNA Affinity Purification (DAP)-Seq dataset [57]. This analysis revealed that 5 out of 7 groups were enriched in targets of known TFs. Of the 7 groups, 4 had enrichment in both annotated function and TF binding. Interestingly, the group that was enriched in transcripts annotated with “negative regulation of ethylene signaling” was also enriched in targets of the ethylene signaling master transcriptional regulator ETHYLENE INSENSITIVE 3 (EIN3), which supported our hypothesis that PaLD could reveal structure that represented biologically-supported subnetworks.

**Table 2.**
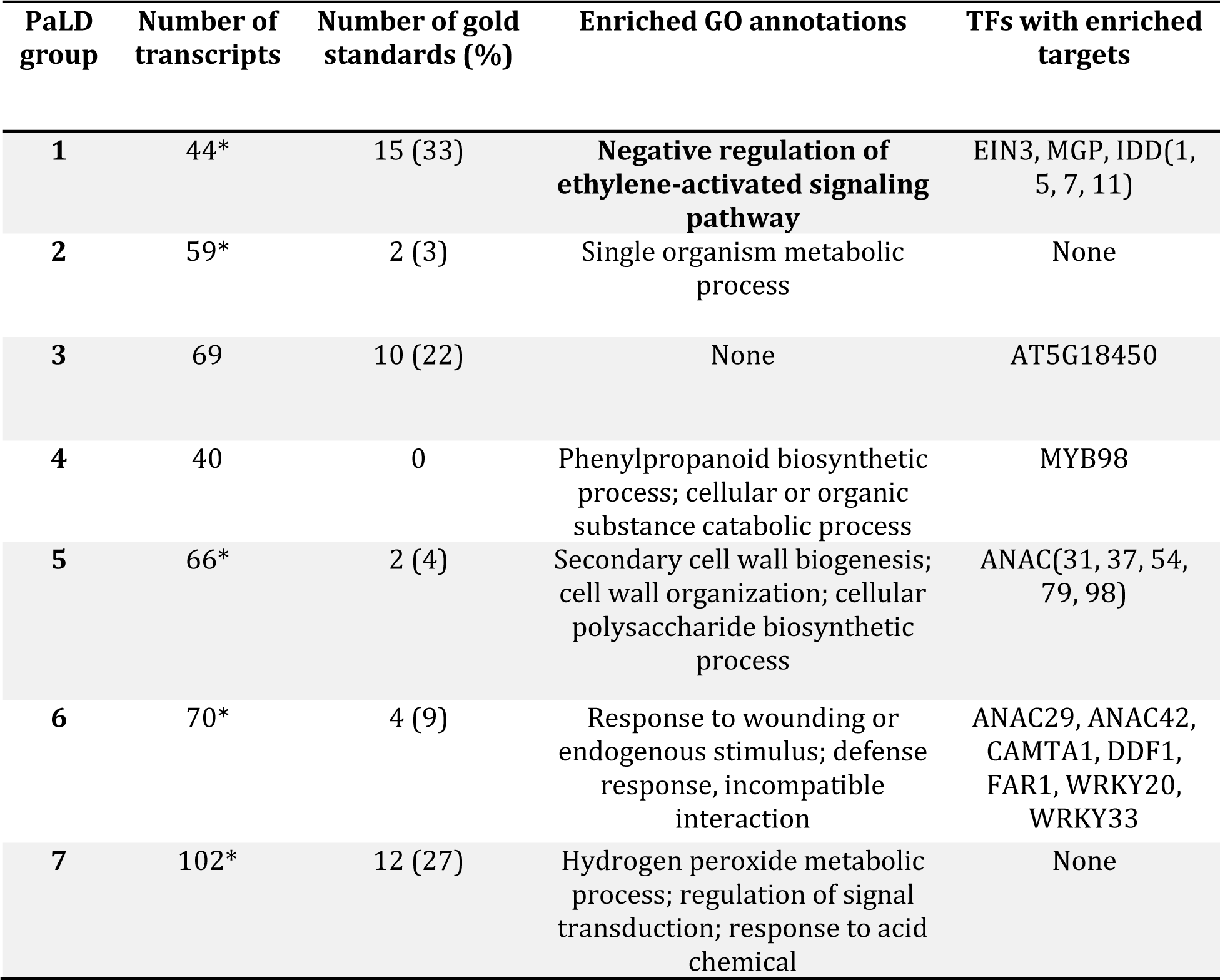
Summary of enrichment results for the PaLD groups generated from the ACC microarray dataset [50]. The 3 GO annotations with the lowest FDRs were reported (adjusted p-value ≤ 0.05). TFs with targets enriched logFC ≥1 relative to the genome were reported. *Indicates groups whose size of some groups is larger in the enrichment analysis due to microarray probes recognizing transcripts from more than one gene.

To further confirm that PaLD could effectively define groups of genes that functioned in the same biological processes, we compared the PaLD groups to a set of core ethylene response genes, called the *gold standards*, which were identified in a meta-analysis of root ethylene responses examining a single time point after treatments that elevated levels of ethylene in roots across three transcriptional datasets [62]. This analysis is shown in the supplemental data, which revealed the patterns of response of this conserved set of ethylene regulated transcripts (Fig. S4). We noted in Table 2 how those consistently ethylene regulated transcripts partitioned between the PaLD groups.

We also examined functional annotation and TF binding in the *k-*means clusters to ask whether these groups were similarly enriched in function and regulation (Table 3). In contrast, 7 out of 10 *k-*means clusters were enriched in annotations, and only 4 out of 10 clusters were enriched in targets of TFs. Clusters A, D, and E highly overlapped with the PaLD groups, while those without enriched TF regulators were those with less overlap with the PaLD groups. The fourth cluster enriched in TF targets was cluster I, which showed a 56% overlap with PaLD group 4, resulting in enrichment in binding sites for MYB98 in both. Only cluster D was enriched in a functional annotation linked to ethylene signaling, and that was highly conserved (67%) with group 1. These results suggested that for this microarray dataset, PaLD (used in combination with a community detection algorithm to reveal subnetworks of transcripts with common temporal response) could reveal biologically meaningful subnetworks, alongside global and local network structure.

**Table 3.** Summary of enrichment results for the top 10 k-means clusters defined in Harkey et al., 2018. The top three GO annotations with the lowest FDRs were reported (adjusted p-value ≤ 0.05). TFs with targets enriched logFC ≥1 relative to the genome were reported. *Indicates groups whose size is larger in the enrichment analysis due to microarray probes recognizing transcripts from more than one gene or smaller due to probes that were not associated with a gene identifier being removed.

| **k-means cluster** | **Number of transcripts** | **Number of gold standards (%)** | **Enriched GO annotations** | **TFs with enriched targets** |
| --- | --- | --- | --- | --- |
| **A** | 115* | 12 (10) | None | FAR1, IDD11, JKD |
| **B** | 50* | 5 (10) | None | None |
| **C** | 41* | 0 | Cellular or organic substance catabolic process | None |
| **D** | 30 | 16 (53) | **Ethylene-activated signaling pathway;** regulation of signal transduction; response to inorganic substance | AT3G10580, AT3G42860, HAT2, MYB3R4, MYB(73, 81, 119), WRKY(20, 31, 33, 40, 55) |
| **E** | 29* | 0 | Response to other organism | CAMTA1, CAMTA5, CBF3 |
| **F** | 21* | 6 (29) | Phenylpropanoid metabolic process | None |
| **G** | 23* | 4 (17) | Cellular component assembly | None |
| **H** | 18 | 0 | None | None |
| **I** | 18 | 0 | Secondary cell wall biogenesis; cellular polysaccharide biosynthetic process; cell wall organization | ANAC45, ANAC96, MYB98, MYB118 |
| **J** | 14* | 1 (1) | None | None |

#### PaLD analysis of transcripts with altered expression in response to elevated auxin

We also examined a second microarray dataset that was simultaneously generated with the ACC dataset described above, which generated a larger group of DE transcripts. PaLD was used to generate a network using the 1246 transcripts whose transcript abundance was significantly changed in roots in response to treatment with the auxin indole-3-acetic acid (IAA). Interestingly, the general structure forms a torus (or doughnut) shape, which is quite different from the “hourglass” shaped ACC network. The Louvain community detection algorithm was applied to this data, yielding 8 groups which are shown in distinct colors in Fig. 5 and are reported in Supplemental Dataset 1. Group 6 consisted of genes that were upregulated most rapidly in response to IAA with a unimodal curve returning to starting levels by the end of 24 hours. Groups 1, 4, and 5 had progressively slower rates of upregulation. Group 2, which was positioned oppositely to groups 1, 4 and 5, also had a unimodal curve, but with decreased transcript abundance in response to auxin, followed by a return to baseline. Fiber plots of each group revealed that the patterns of transcript abundance over time were slightly shifted from one group to the next. We reasoned that these results might be indicative of waves of transcripts that were activated at various times from the start of hormone treatment.

**Fig. 5.**
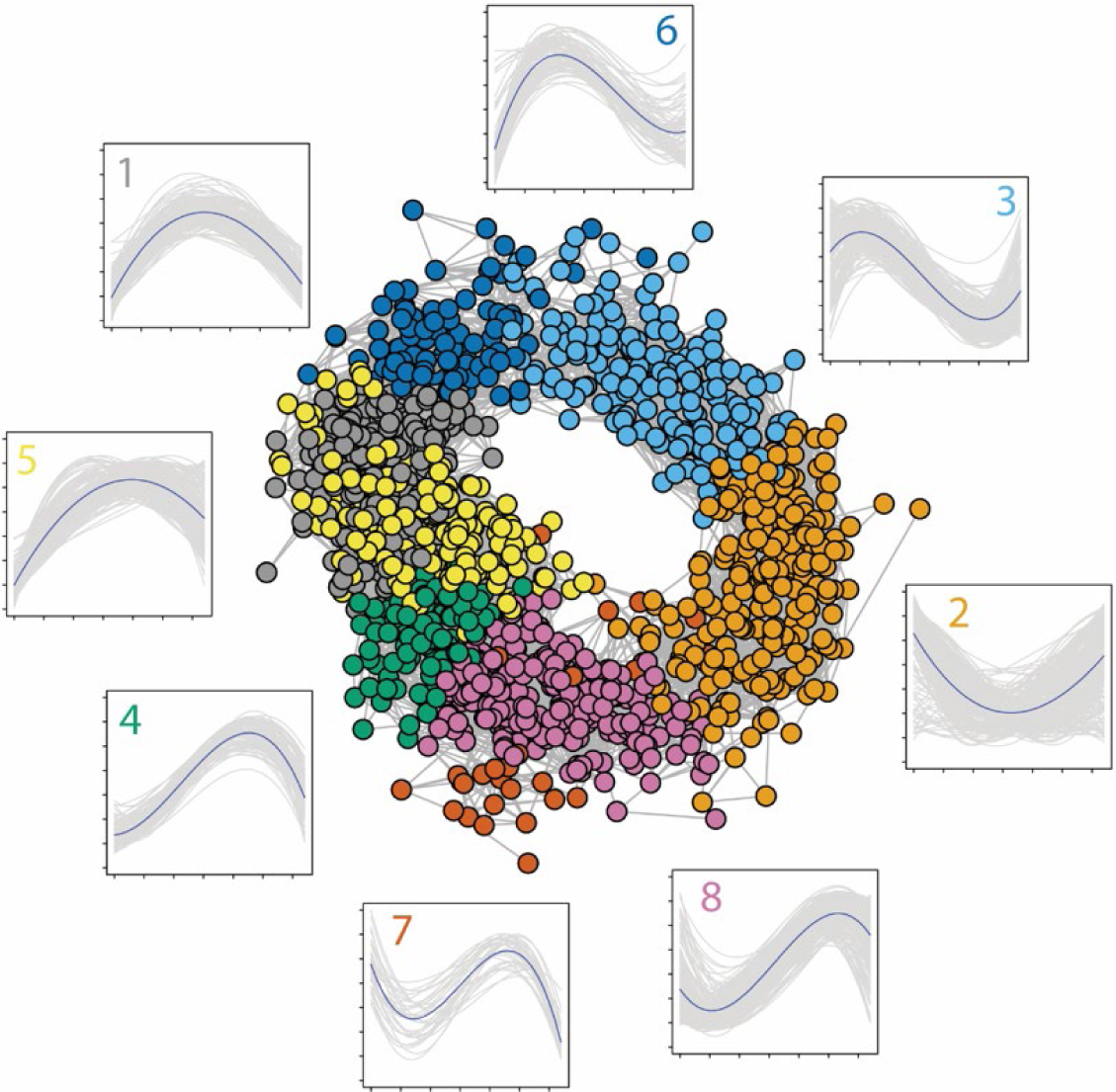
PaLD analysis of the 1246 transcripts with altered transcript abundance in response to treatment with IAA, an auxin [52]. The network diagram shows each PaLD group in a different color. Fiber plots of the PaLD groups illustrate transcript abundance over time with the number of each group matching the color of each group. Data is considered in log time and has been min-max scaled.

To further contrast the differences in network structures between those for the ACC and IAA microarray datasets, we generated videos depicting rotations of three-dimensional renderings of the PaLD networks of each of these structures (Supplemental Movies 1 and 2). From these videos we produced snapshots of the networks from representative angles (Fig. 6). From these images it is clear that many of the groups that appear to overlap with one another in the two-dimensional networks were spatially, and thus temporally, distinct. It is interesting to contrast the local and global structural information provided by PaLD over standard node-labelling methods.

**Fig. 6.**
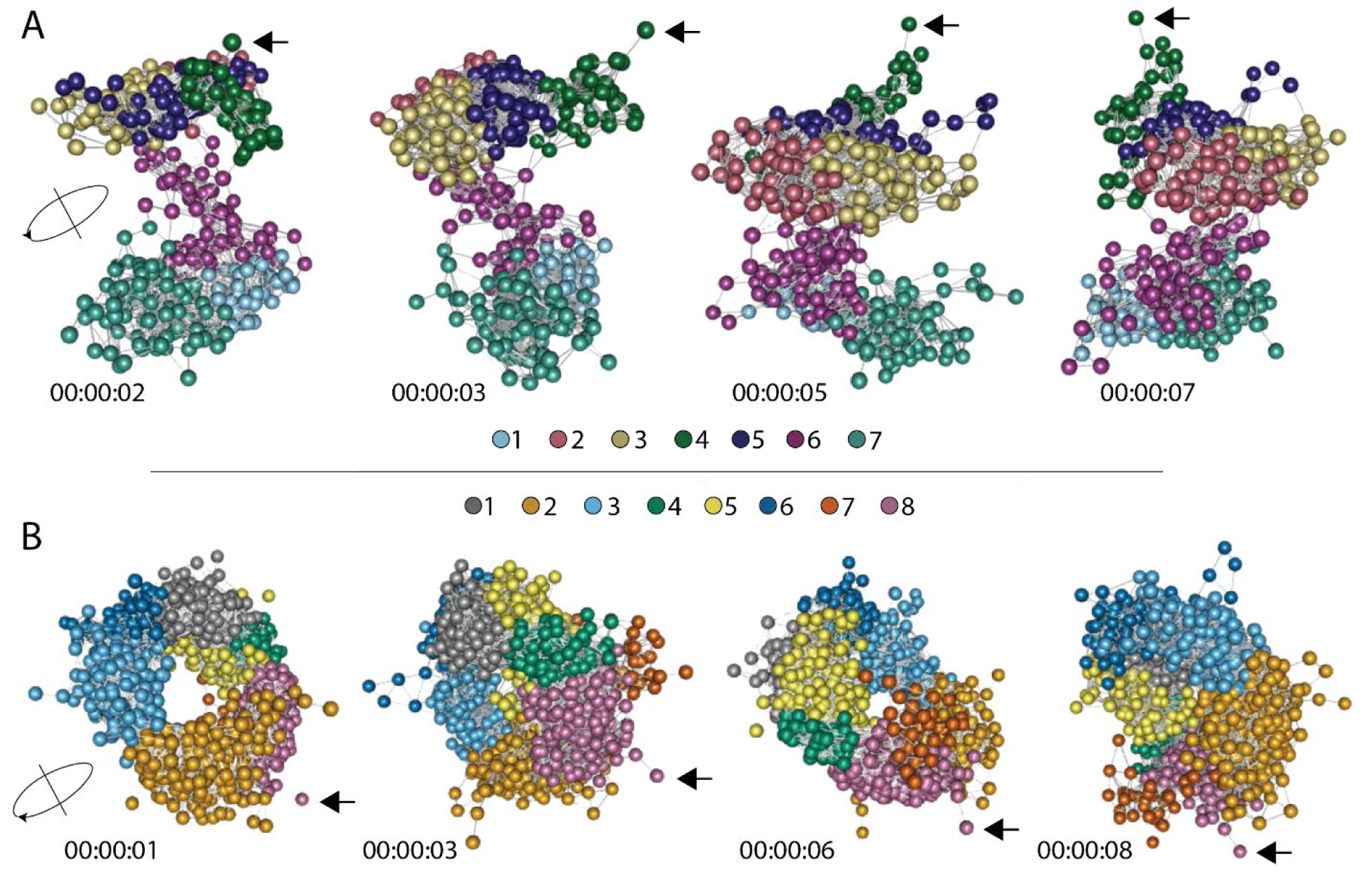
Screenshots from movies representing three-dimensional network structures for the ACC and IAA microarray datasets analyzed using PaLD. (A) The ACC PaLD network is shown from 4 representative angles depicted as rotating counterclockwise around the axis (left) from left to right. Timestamps from movies are included below the frame in which the screenshot was obtained. (B) The IAA PaLD network is shown from 4 representative angles illustrated as rotating clockwise around the axis (left) from left to right. Timestamps from movies are included below the frame in which the screenshot was obtained. In both panels, the colors represent the PaLD groups. Black arrows point to the same gene in each network for frame of reference.

We compared the distribution of transcripts between the PaLD groups and the top ten *k-* means clusters for the IAA dataset, in a matrix (Table S2). The PaLD and *k-*means methods appeared to sort temporally-linked transcripts in different ways, providing a basis for comparing enrichment between the PaLD groups and *k-*means clusters. We looked for enriched function and enriched TF binding to genes in these PaLD groups and compared them to the *k-*means clusters. Enriched GO annotations were found in all 8 PaLD groups and 8 out of 10 (80%) *k-*means clusters (Table 4 and Table S3). On the other hand, 5 out of 8 (63%) and 7 out of 10 (70%) PaLD groups and *k-*means clusters, respectively, were enriched in targets of TFs. As predicted, each set of groups/clusters had groups enriched in auxin signaling, with the PaLD IAA group 5 and 6 and *k-*means cluster G having this annotation. Nevertheless, examination of each of the summary tables revealed a number of differences in the qualitative output. For example, the GO annotation “covalent chromatin modification” was enriched in PaLD group 4 but was not enriched in any of the *k*-means clusters (Table 4 and Table S3). Similarly, genes functioning in “root morphogenesis” and “root meristem growth” were enriched in PaLD group 5 but were not enriched in any of the *k*-means clusters (Table 4 and Table S3). These results highlight the ability of PaLD to reveal new functional information regarding time series data that is biologically relevant.

**Table 4.** Summary of enrichment results for the PaLD groups generated from the IAA microarray dataset [52]. The top three GO annotations with the lowest FDRs were reported (adjusted p-value ≤ 0.05). TFs with targets enriched logFC ≥1 relative to the genome were reported. *This indicates that the group size is smaller than expected due to probes that were not associated with a gene identifier being removed.

| PaLD group | Number of transcripts | Enriched GO annotations | TFs with enriched targets |
| --- | --- | --- | --- |
| 1 | 193* | Ribosomal large subunit biogenesis; ribosome assembly; pyrimidine nucleotide biosynthetic process | TRP1 |
| 2 | 246 | Cell wall organization; response to water deprivation; sulfur compound metabolic process | AT1G01250, BEH3, bZIP51, CAMTA5, FAR1, LEP, MYB96, RAP2.1 |
| 3 | 185* | Glycosyl or organonitrogen compound catabolic process; response to wounding | ERF2, ERF5, RAP2.9, WIND4 |
| 4 | 101* | Cell cycle; covalent chromatin modification; regulation of protein modification process | None |
| 5 | 193 | Auxin-activated signaling pathway; root morphogenesis; root meristem growth | None |
| 6 | 94 | **Cellular response to auxin stimulus** | None |
| 7 | 32 | Cellular lipid metabolic process | ANAC(7,37,38,45,50,53, 54,57,79,87,92,96), AT4G26030, MYB(49,74,80,99,107) |
| 8 | 228* | Cell division; **hormone-mediated signaling pathway**; response to abiotic stimulus | DDF2, DOF4.2, JKD, LBD23, LOB, MYB3R4 |

To demonstrate the application of PaLD in the analysis of time series gene expression data derived using RNA-seq, we applied this same analysis pipeline to two additional datasets that examined gene expression over time in response to ethylene gas [51] (Fig. S5 and Table S4) or salt stress [53] (Fig. S6 and Tables S5-S7). For more details on the results of these analyses, see Supplemental Materials.

#### PaLD neighborhoods revealed connections between Auxin Response Factor 19 and root responses in the IAA microarray dataset

We asked whether PaLD could reveal smaller subnetworks of interest in the IAA microarray dataset. We used a second approach to look for neighborhoods of like-responding genes in the PaLD network, by generating lists of nearest neighbors for each gene. It is important to note that neighborhoods can be of varying size reflecting heterogeneity in the data. To illustrate this approach, we selected the rapidly upregulated gene encoding Auxin Response Factor 19 (ARF19), which drives auxin gene regulatory networks in roots. The *ARF19* neighborhood was highlighted on the PaLD network (Fig. 7A) and the temporal responses of the genes in this neighborhood were illustrated in the fiber plot in Fig. 7B and are listed in Supplemental Data Set 1.

**Fig.7.**
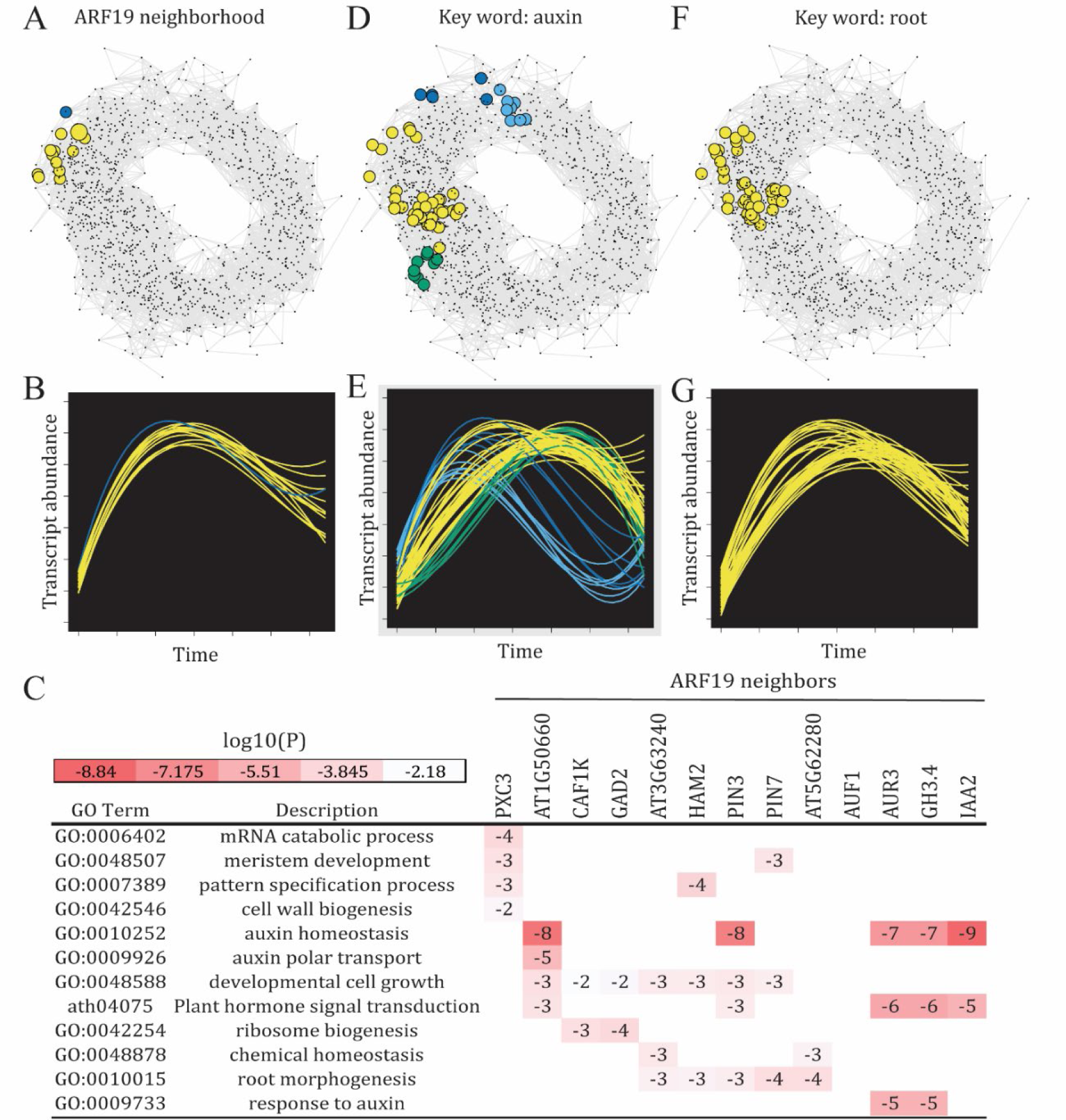
Functionally relevant neighborhoods in the IAA network. (A) On the top, the neighborhood of Auxin Response Factor 19 (ARF19), with each node colored according to PaLD groups as displayed in Fig. 5. (B) This fiber plot shows transcript abundance over time for all transcripts in the ARF19 neighborhood. Data is considered in log time and has been min-max scaled. (C) The enrichment output for the neighborhoods of the neighbors of ARF19. The names of the genes or locus identifier of ARF19 neighbors are shown along the top of the heatmap. Values indicate the p values in log base 10 rounded to the nearest whole number for enrichment of that term in each gene’s neighborhood relative to the whole genome. Neighborhoods that were not enriched in a GO term do not have a value shown. (D and F) Neighborhoods for each transcript were queried for conserved function using Metascape and transcripts annotated with the GO term “auxin” or “root” are highlighted on the network with colors corresponding to PaLD groups as displayed in Fig. 5. (E and G) The fiber plots for these annotated groups are shown with the same color coding.

We next asked whether PaLD could resolve functional differences in subnetworks with high degrees of overlap (e.g., neighborhoods of the genes directly connected to *ARF19*). Interestingly, most of the genes directly connected to *ARF19* in the network encoded proteins known to be involved in auxin signaling and transport, including PINFORMED 3 (PIN3), PIN7, and AUXIN-INDUCIBLE 2 (IAA2). Fig. 7C shows the enriched biological functions for each of the neighborhoods of the genes directly connected to *ARF19* in the PaLD network, with many neighborhoods sharing enriched GO annotations with one another and many neighborhoods enriched in different GO annotations. For example, the neighborhoods of 2 auxin transporters, PIN3 and PIN7, were both enriched in GO terms linked to developmental cell growth and root morphogenesis, while only the neighborhood of PIN3 was enriched in auxin homeostasis and plant hormone signal transduction and only PIN7 in meristem development (Fig 7C).

We also sought to reveal subnetworks by searching for specific GO annotations (a top-down approach). For contrast, here, we employed a different enrichment analysis software, Metascape [67], which employs a greater number of GO knowledgebases compared to AgriGO. We automated a process using Metascape to identify all the GO terms enriched in each of the 1246 neighborhoods in the IAA network. We then used this data to search for neighborhoods enriched in GO terms that included “auxin” or “root” and highlighted these neighborhoods on the PaLD network in Fig. 7D and F, respectively. The transcripts involved in root-related processes formed a localized niche in the network (Fig. 7F), which overlapped significantly with the *ARF19* neighborhood, while transcripts associated with auxin-related processes were more dispersed throughout a larger region of the network (Fig. 7D). Likewise, the fiber plot of the auxin-associated neighbors was revealed to be variable (Fig. 7E), while the fiber plot of the root-associated transcripts was more consistent in temporal responses (Fig. 7G). Taken together, these findings revealed that PaLD neighborhoods could be used to predict subnetworks, and that a top-down approach could be employed to search for these networks.

## Discussion

A central challenge in systems biology is the inference of gene relationships from patterns found in transcriptomic data. In particular, time series transcript abundance data include a temporal dimension that allows us to make predictions about like-responding transcripts, due to common transcriptional controls or biological function of the proteins that they encode. These transcriptomic datasets can be used to address questions including how hormones or physiological stress alter the transcriptional landscape in plants. Such experiments reveal the set of genes that have altered expression under elevated levels of plant hormones or salinity, for instance, providing new information about the pathways and mechanisms that are activated in response to these factors. In this paper, we analyzed several time series transcriptomic datasets, generated for the model plant Arabidopsis, to delve more deeply into the mechanisms driving these complex regulatory processes, by applying new methods to query the data for predictive patterns. The datasets used in this analysis were selected to focus on hormone and stress response gene regulatory networks in Arabidopsis.

The wide variety of methods which can be employed in bioinformatics can make it challenging to determine which methods to apply and, beyond this, may require the user to make decisions that influence the results. Specifically, widely employed methods such as *k-* means and hierarchical clustering require the user to provide additional inputs (e.g., the number of clusters) or select values from a parameter space (e.g., to minimize some cost function). Many methods used to cluster time series data provide limited information about the global structure of the data, lacking a way to visualize how transcripts are both directly and indirectly connected to one another within a larger network. Dimension reduction techniques, such as principal component analysis, are often utilized to provide a flat 2-dimensional display of the data. Since such methods can compress valuable structural features, network*-*based methods can also be used to reveal informative structure. Yet, again, methods for creating such networks may force the user to select parameters (e.g., number of neighbors, or a threshold for the maximum distance between neighbors) and these choices can influence the results obtained. In light of these challenges, the need to share straightforward modeling algorithms for holistically reflecting local and global structure (including delineated groups or clusters), which can be easily applied to biological datasets, is more critical than ever before.

In this paper, we present the first application of Partitioned Local Depth (PaLD) to time series transcript abundance data and provide a self-contained description of the approach. Our focus here is on the shape of the curve for transcript abundance over time; we measured similarity in transcript response using fitted cubic polynomials to expression levels considered in log-time to reflect the rise/fall behavior typical of time series transcriptomic data; other application-dependent choices are possible and can be used to best fit the response patterns of individual datasets. The networks defined by PaLD provide a natural set of temporally-linked transcripts, defined by the endogenous patterns of the data. The method provides a transparent way to construct a network which does not require additional user input parameters. The recently introduced method was defined to reflect social interactions, and thus the neighbors and overall structure revealed by PaLD can be intuitively interpreted. For further discussion, see [46]. An R package for PaLD with detailed examples describing ways that it can be employed can be found at https://github.com/moorekatherine/pald (Katherine Moore, Kenneth Berenhaut, and Lucy D’Agostino McGowan, Github Package, 2022).

To contextualize the results and illustrate the value of this new method, we analyzed four time series transcriptomic datasets that were generated in Arabidopsis. Three of the datasets examined hormone response [17–19], while the other examined the stress responses to elevated salt [53]. PaLD analysis of a microarray dataset of root samples treated with ACC, a precursor to the hormone ethylene [50], revealed proportionally more groups with enrichment in functional annotations and targets of TFs as compared to a previously employed *k-*means clustering method. Most striking was the significant enrichment in one of the PaLD groups in ethylene signaling. This group was enriched in targets of the ethylene signaling master transcriptional regulator ETHYLENE INSENSITIVE 3 (EIN3). In comparison, the *k-*means method also yielded a group that was enriched in the function “ethylene signaling,” but this group was not enriched in targets of EIN3. The application of PaLD to an ethylene gas RNA-Seq dataset [51] also resulted in two groups enriched in ethylene signaling, with one of these groups enriched in targets of EIN3.

Consistent with the ability of ACC to remodel root architecture [62,63], PaLD and *k-*means clustering methods revealed groups that were enriched in secondary cell wall biogenesis, which is necessary for driving primary and lateral root growth and root hair formation. The PaLD groups and *k-*means clusters with this annotation also showed enrichment in targets of the NAC protein family of TFs, suggesting that the NAC family may drive the gene regulatory networks remodeling root growth under conditions of elevated ethylene.

We also applied the PaLD method to analyze a microarray dataset in which roots were treated with IAA. Both the PaLD and *k-*means clustering methods resulted in similar numbers of groups that were enriched in functional annotations and targets of TFs but differ in distribution of genes between these groups and thus the content of these enrichments. A benefit of using the PaLD approach is that the number of like-responding transcripts can vary across the set and, in contrast to common nearest-neighbor methods, is not a single quantity that must be specified by the user. This allows us to identify meaningful sets of temporally similar transcripts that are directly linked to given genes of interest. For illustrative purposes, we examined the neighborhood of *Auxin Response Factor 19* (*ARF19*) revealing transcripts that were directly connected to *ARF19* included genes encoding the auxin transporters PIN3 and PIN7. This finding is consistent with these transcripts having similar kinetic responses to auxin, which may be due to ARF19 driving PIN3 and PIN7 transcription or with all 3 transcripts being controlled by a common TF. We also found significant overlap between the ARF19 neighborhood and transcripts that were associated with GO annotations linked to “root.” These results review the utility of the localized neighborhoods that PaLD generates as a way to reveal transcriptional networks.

The PaLD approach provides network models visualizations that are rich and informative spatial representations of the relationships among temporally-linked transcripts. The resulting local and global structure provides insight into the relationships between genes. We showed how PaLD can be used to model biological networks and reveal groups of genes with enriched function and regulation, which can then be employed to predict subnetworks supported by functional and TF-target information. The PaLD networks can also be employed to obtain a collection of neighboring genes, a feature which may be valuable in exploratory analysis. Ultimately, the unique individual gene perspective offers a plethora of opportunities for exploration into the functional and regulatory patterns of different subnetworks. Finally, this method used in combination with community detection algorithms, such as the Louvain method, can result in informative groups of transcripts. These groups can be examined for enriched functional annotation and transcriptional regulation which can generate hypotheses about gene interactions and biological implications that can then be tested experimentally. In particular, Arabidopsis T-DNA insertional mutants are widely available to test which genes might be misregulated downstream of knockout mutant in a TF, thereby revealing transcriptional networks.

Taken together, the PaLD approach allows time series transcriptomic data, which is often rich and complex, to become more interpretable and manageable in a research setting. In light of the increased specialization of researchers across the sciences, the need to create straightforward modeling algorithms that can be easily applied to datasets by biologists with limited training in bioinformatics is more critical than ever before. Ultimately, we demonstrated here that PaLD is a straightforward method that can be used to process transcriptomic data to reveal enrichment in function and regulation, which is the first step in modeling complex biological processes that can then be validated experimentally in the laboratory.

## Author contributions

MGK: Formal analysis, Investigation, Writing-Original Draft Preparation and Review and Editing; KSB: Conceptualization, Formal analysis, Writing-Original Draft Preparation and Review and Editing; KEM: Formal analysis, Investigation, Writing-Original Draft Preparation; EEA: Formal analysis, Writing-Original Draft Preparation and Review and Editing, AFH: Investigation, Writing-Review and Editing, JKM: Investigation, Writing-Review and Editing, CNC: Formal Analysis, JX: Formal Analysis, SSJ: Investigation; DJJ: Formal Analysis, Writing-Review and Editing; JLN: Formal Analysis, Writing-Review and Editing, GKM: Conceptualization, Project Administration, Funding Acquisition, and Writing-Review and Editing.

## References

1. Hemonnot-Girard A-L, Meersseman C, Pastore M, Garcia V, Linck N, Rey C, et al. Comparative analysis of transcriptome remodeling in plaque-associated and plaque-distant microglia during amyloid-β pathology progression in mice. Journal of Neuroinflammation. 2022;19: 234. doi:10.1186/s12974-022-02581-0

2. Li Y, Shah-Simpson S, Okrah K, Belew AT, Choi J, Caradonna KL, et al. Transcriptome Remodeling in Trypanosoma cruzi and Human Cells during Intracellular Infection. PLOS Pathogens. 2016;12: e1005511. doi:10.1371/journal.ppat.1005511

3. Wright Meredith S., Jacobs Michael R., Bonomo Robert A., Adams Mark D. Transcriptome Remodeling of Acinetobacter baumannii during Infection and Treatment. mBio. 2017;8: e02193–16. doi:10.1128/mBio.02193-16

4. Wilkinson D, Maršíková J, Hlaváček O, Gilfillan GD, Ježková E, Aaløkken R, et al. Transcriptome Remodeling of Differentiated Cells during Chronological Ageing of Yeast Colonies: New Insights into Metabolic Differentiation. Thevissen K, editor. Oxidative Medicine and Cellular Longevity. 2018;2018: 4932905. doi:10.1155/2018/4932905

5. Landa M, Burns AS, Roth SJ, Moran MA. Bacterial transcriptome remodeling during sequential co-culture with a marine dinoflagellate and diatom. The ISME Journal. 2017;11: 2677–2690. doi:10.1038/ismej.2017.117

6. Pawełczyk J, Brzostek A, Minias A, Płociński P, Rumijowska-Galewicz A, Strapagiel D, et al. Cholesterol-dependent transcriptome remodeling reveals new insight into the contribution of cholesterol to Mycobacterium tuberculosis pathogenesis. Scientific Reports. 2021;11: 12396. doi:10.1038/s41598-021-91812-0

7. Mazille M, Buczak K, Scheiffele P, Mauger O. Stimulus-specific remodeling of the neuronal transcriptome through nuclear intron-retaining transcripts. The EMBO Journal. 2022;41: e110192. doi:10.15252/embj.2021110192

8. Szenajch J, Szabelska-Beręsewicz A, Świercz A, Zyprych-Walczak J, Siatkowski I, Góralski M, et al. Transcriptome Remodeling in Gradual Development of Inverse Resistance between Paclitaxel and Cisplatin in Ovarian Cancer Cells. International Journal of Molecular Sciences. 2020;21. doi:10.3390/ijms21239218

9. Mozejko-Ciesielska J, Pokoj T, Ciesielski S. Transcriptome remodeling of Pseudomonas putida KT2440 during mcl-PHAs synthesis: effect of different carbon sources and response to nitrogen stress. Journal of Industrial Microbiology and Biotechnology. 2018;45: 433–446. doi:10.1007/s10295-018-2042-4

10. Schreiner WP, Pagliuso DC, Garrigues JM, Chen JS, Aalto AP, Pasquinelli AE. Remodeling of the Caenorhabditis elegans non-coding RNA transcriptome by heat shock. Nucleic Acids Research. 2019;47: 9829–9841. doi:10.1093/nar/gkz693

11. Sima C, Hua J, Jung S. Inference of Gene Regulatory Networks Using Time-Series Data: A Survey. Current Genomics. 2009;10: 416–429. doi:10.2174/138920209789177610

12. Yeung KY, Dombek KM, Lo K, Mittler JE, Zhu J, Schadt EE, et al. Construction of regulatory networks using expression time-series data of a genotyped population. Proceedings of the National Academy of Sciences. 2011;108: 19436–19441. doi:10.1073/pnas.1116442108

13. Ding J, Bar-Joseph Z. Analysis of time-series regulatory networks. Current Opinion in Systems Biology. 2020;21: 16–24. doi:10.1016/j.coisb.2020.07.005

14. Kimura S, Fukutomi R, Tokuhisa M, Okada M. Inference of Genetic Networks From Time-Series and Static Gene Expression Data: Combining a Random-Forest-Based Inference Method With Feature Selection Methods. Frontiers in Genetics. 2020;11. Available: https://www.frontiersin.org/articles/10.3389/fgene.2020.595912

15. Zhang Y, Chang X, Liu X. Inference of gene regulatory networks using pseudo-time series data. Bioinformatics. 2021;37: 2423–2431. doi:10.1093/bioinformatics/btab099

16. Lu J, Dumitrascu B, McDowell IC, Jo B, Barrera A, Hong LK, et al. Causal network inference from gene transcriptional time-series response to glucocorticoids. PLOS Computational Biology. 2021;17: e1008223. doi:10.1371/journal.pcbi.1008223

17. Ovens K, Eames BF, McQuillan I. Comparative Analyses of Gene Co-expression Networks: Implementations and Applications in the Study of Evolution. Frontiers in Genetics. 2021;12. Available: https://www.frontiersin.org/articles/10.3389/fgene.2021.695399

18. Montenegro JD. Gene Co-expression Network Analysis. In: Edwards D, editor. Plant Bioinformatics: Methods and Protocols. New York, NY: Springer US; 2022. pp. 387–404. doi:10.1007/978-1-0716-2067-0_19

19. Yin W, Mendoza L, Monzon-Sandoval J, Urrutia AO, Gutierrez H. Emergence of co-expression in gene regulatory networks. PLOS ONE. 2021;16: e0247671. doi:10.1371/journal.pone.0247671

20. Paci P, Fiscon G, Conte F, Wang R-S, Farina L, Loscalzo J. Gene co-expression in the interactome: moving from correlation toward causation via an integrated approach to disease module discovery. npj Systems Biology and Applications. 2021;7: 3. doi:10.1038/s41540-020-00168-0

21. Kolberg L, Kerimov N, Peterson H, Alasoo K. Co-expression analysis reveals interpretable gene modules controlled by trans-acting genetic variants. Parker SC, Wittkopp PJ, Parker SC, Ruffieux H, editors. eLife. 2020;9: e58705. doi:10.7554/eLife.58705

22. Zainal-Abidin R-A, Harun S, Vengatharajuloo V, Tamizi A-A, Samsulrizal NH. Gene Co-Expression Network Tools and Databases for Crop Improvement. Plants. 2022;11. doi:10.3390/plants11131625

23. E. E. Allen, J. L. Norris, D. J. John, S. J. Thomas, W. H. Turkett Jr., J. S. Fetrow. Comparison of Co-temporal Modeling Algorithms on Sparse Experimental Time Series Data Sets. 2010 IEEE International Conference on BioInformatics and BioEngineering. 2010. pp. 79–85. doi:10.1109/BIBE.2010.21

24. Rockne RC, Branciamore S, Qi J, Frankhouser DE, O’Meally D, Hua W-K, et al. State-Transition Analysis of Time-Sequential Gene Expression Identifies Critical Points That Predict Development of Acute Myeloid Leukemia. Cancer Research. 2020;80: 3157–3169. doi:10.1158/0008-5472.CAN-20-0354

25. Lee J, Heath LS, Grene R, Li S. Comparing time series transcriptome data between plants using a network module finding algorithm. Plant Methods. 2019;15: 61. doi:10.1186/s13007-019-0440-x

26. Spies D, Ciaudo C. Dynamics in Transcriptomics: Advancements in RNA-seq Time Course and Downstream Analysis. Computational and Structural Biotechnology Journal. 2015;13: 469–477. doi:10.1016/j.csbj.2015.08.004

27. Wu Y, Xue L, Huang W, Deng M, Lin Y. Profiling transcription factor activity dynamics using intronic reads in time-series transcriptome data. PLOS Computational Biology. 2022;18: e1009762. doi:10.1371/journal.pcbi.1009762

28. Petricka JJ, Winter CM, Benfey PN. Control of Arabidopsis Root Development. Annu Rev Plant Biol. 2012;63: 563–590. doi:10.1146/annurev-arplant-042811-105501

29. Niklison-Chirou MV, Agostini M, Amelio I, Melino G. Regulation of Adult Neurogenesis in Mammalian Brain. International Journal of Molecular Sciences. 2020;21. doi:10.3390/ijms21144869

30. Yosef N, Regev A. Impulse Control: Temporal Dynamics in Gene Transcription. Cell. 2011;144: 886–896. doi:10.1016/j.cell.2011.02.015

31. de Luis Balaguer MA, Fisher AP, Clark NM, Fernandez-Espinosa MG, Möller BK, Weijers D, et al. Predicting gene regulatory networks by combining spatial and temporal gene expression data in *Arabidopsis* root stem cells. Proc Natl Acad Sci USA. 2017;114. doi:10.1073/pnas.1707566114

32. Mochida K, Koda S, Inoue K, Nishii R. Statistical and Machine Learning Approaches to Predict Gene Regulatory Networks From Transcriptome Datasets. Front Plant Sci. 2018;9: 1770–1770. doi:10.3389/fpls.2018.01770

33. Omranian N, Eloundou-Mbebi JMO, Mueller-Roeber B, Nikoloski Z. Gene regulatory network inference using fused LASSO on multiple data sets. Sci Rep. 2016;6: 20533–20533. doi:10.1038/srep20533

34. Van den Broeck L, Gordon M, Inzé D, Williams C, Sozzani R. Gene Regulatory Network Inference: Connecting Plant Biology and Mathematical Modeling. Frontiers in Genetics. 2020;11. Available: https://www.frontiersin.org/article/10.3389/fgene.2020.00457

35. Koch CM, Chiu SF, Akbarpour M, Bharat A, Ridge KM, Bartom ET, et al. A Beginner’s Guide to Analysis of RNA Sequencing Data. Am J Respir Cell Mol Biol. 2018;59: 145–157. doi:10.1165/rcmb.2017-0430TR

36. Altman N, Krzywinski M. Clustering. Nature Methods. 2017;14: 545–546. doi:10.1038/nmeth.4299

37. Jamail I, Moussa A. Current State-of-the-Art of Clustering Methods for Gene Expression Data with RNA-Seq. Applications of Pattern Recognition. IntechOpen; 2020. doi:10.5772/intechopen.94069

38. Everitt BS, Landau S, Leese M, Stahl D. Cluster Analysis: Everitt/Cluster Analysis. Chichester, UK: John Wiley & Sons, Ltd; 2011. doi:10.1002/9780470977811

39. Kaufman L, Rousseeuw PJ. Finding Groups in Data: an Introduction to Cluster Analysis. Hoboken: John Wiley & Sons, Inc.; 2009. Available: http://www.SLQ.eblib.com.au/patron/FullRecord.aspx?p=469065

40. von Luxburg U. A tutorial on spectral clustering. Stat Comput. 2007;17: 395–416. doi:10.1007/s11222-007-9033-z

41. Ernst J, Bar-Joseph Z. STEM: a tool for the analysis of short time series gene expression data. BMC Bioinformatics. 2006;7: 191. doi:10.1186/1471-2105-7-191

42. Ernst J, Vainas O, Harbison CT, Simon I, Bar-Joseph Z. Reconstructing dynamic regulatory maps. Molecular Systems Biology. 2007;3: 74. doi:10.1038/msb4100115

43. Oh V-KS, Li RW. Temporal Dynamic Methods for Bulk RNA-Seq Time Series Data. Genes. 2021;12. doi:10.3390/genes12030352

44. Y. Asano, T. Ogawa, S. Shichino, S. Ueha, K. Matsushima, A. Ogura. Time-Series Analysis of Gene Correlation Networks based on Single-Cell Transcriptome Data. 2021 IEEE International Conference on Bioinformatics and Biomedicine (BIBM). 2021. pp. 2134–2141. doi:10.1109/BIBM52615.2021.9669412

45. Jung I, Jo K, Kang H, Ahn H, Yu Y, Kim S. TimesVector: a vectorized clustering approach to the analysis of time series transcriptome data from multiple phenotypes. Bioinformatics. 2017;33: 3827–3835. doi:10.1093/bioinformatics/btw780

46. Berenhaut KS, Moore KE, Melvin RL. A social perspective on perceived distances reveals deep community structure. Proceedings of the National Academy of Sciences. 2022;119: e2003634119. doi:10.1073/pnas.2003634119

47. A. M. Mehar, K. Matawie, A. Maeder. Determining an optimal value of K in K-means clustering. 2013 IEEE International Conference on Bioinformatics and Biomedicine. 2013. pp. 51–55. doi:10.1109/BIBM.2013.6732734

48. Blondel VD, Guillaume J-L, Lambiotte R, Lefebvre E. Fast unfolding of communities in large networks. J Stat Mech. 2008;2008: P10008. doi:10.1088/1742-5468/2008/10/P10008

49. Newman MEJ. Communities, modules and large-scale structure in networks. Nature Phys. 2012;8: 25–31. doi:10.1038/nphys2162

50. Harkey AF, Watkins JM, Olex AL, DiNapoli KT, Lewis DR, Fetrow JS, et al. Identification of Transcriptional and Receptor Networks That Control Root Responses to Ethylene. Plant Physiology. 2018;176: 2095–2118. doi:10.1104/pp.17.00907

51. Chang KN, Zhong S, Weirauch MT, Hon G, Pelizzola M, Li H, et al. Temporal transcriptional response to ethylene gas drives growth hormone cross-regulation in Arabidopsis. Weigel D, editor. eLife. 2013;2: e00675. doi:10.7554/eLife.00675

52. Lewis DR, Olex AL, Lundy SR, Turkett WH, Fetrow JS, Muday GK. A Kinetic Analysis of the Auxin Transcriptome Reveals Cell Wall Remodeling Proteins That Modulate Lateral Root Development in Arabidopsis. The Plant Cell. 2013;25: 3329–3346. doi:10.1105/tpc.113.114868

53. Wu T-Y, Goh H, Azodi CB, Krishnamoorthi S, Liu M-J, Urano D. Evolutionarily conserved hierarchical gene regulatory networks for plant salt stress response. Nature Plants. 2021;7: 787–799. doi:10.1038/s41477-021-00929-7

54. Gamalero E, Glick BR. Recent Advances in Bacterial Amelioration of Plant Drought and Salt Stress. Biology (Basel). 2022;11: 437. doi:10.3390/biology11030437

55. Chen H, Bullock DA, Alonso JM, Stepanova AN. To Fight or to Grow: The Balancing Role of Ethylene in Plant Abiotic Stress Responses. Plants. 2022;11. doi:10.3390/plants11010033

56. Kamada T, Kawai S. An algorithm for drawing general undirected graphs. Information Processing Letters. 1989;31: 7–15. doi:10.1016/0020-0190(89)90102-6

57. O’Malley RC, Huang S-SC, Song L, Lewsey MG, Bartlett A, Nery JR, et al. Cistrome and Epicistrome Features Shape the Regulatory DNA Landscape. Cell. 2016;165: 1280– 1292. doi:10.1016/j.cell.2016.04.038

58. Harkey AF, Sims KN, Muday GK. A new tool for discovering transcriptional regulators of co-expressed genes predicts gene regulatory networks that mediate ethylene-controlled root development. in silico Plants. 2020;2: diaa006. doi:10.1093/insilicoplants/diaa006

59. Tian T, Liu Y, Yan H, You Q, Yi X, Du Z, et al. agriGO v2.0: a GO analysis toolkit for the agricultural community, 2017 update. Nucleic Acids Research. 2017;45: W122–W129. doi:10.1093/nar/gkx382

60. Zhou Y, Zhou B, Pache L, Chang M, Khodabakhshi AH, Tanaseichuk O, et al. Metascape provides a biologist-oriented resource for the analysis of systems-level datasets. Nature Communications. 2019;10: 1523. doi:10.1038/s41467-019-09234-6

61. Andrews DF. Plots of High-Dimensional Data. Biometrics. 1972;28: 125–136. doi:10.2307/2528964

62. Harkey AF, Yoon GM, Seo DH, DeLong A, Muday GK. Light Modulates Ethylene Synthesis, Signaling, and Downstream Transcriptional Networks to Control Plant Development. Front Plant Sci. 2019;10: 1094. doi:10.3389/fpls.2019.01094

63. Muday GK, Rahman A, Binder BM. Auxin and ethylene: collaborators or competitors? Trends in Plant Science. 2012;17: 181–195. doi:10.1016/j.tplants.2012.02.001

64. Love MI, Huber W, Anders S. Moderated estimation of fold change and dispersion for RNA-seq data with DESeq2. Genome Biology. 2014;15: 550. doi:10.1186/s13059-014-0550-8

