## Supplemental Figures for "Partitioned Local Depth analysis of time series transcript abundance data reveals elaborate community structure"

### Supplemental Materials

#### Supplemental Methods

##### Microarray datasets:

Microarray datasets generated in parallel and published in [52] ("auxin dataset") and [50] ("ACC dataset"), were used in this study. Arabidopsis seedlings at five days after germination were transferred to control growth media or media containing either 1-amino-cyclopropane-1-carboxylic acid (ACC, 1 μM), which is a precursor to ethylene, or indole-3-acetic acid (IAA, 1 μM), which is an auxin. RNA was isolated from roots at eight time points: 0, 0.5, 1, 2, 4, 8, 12, and 24 hours. Triplicates of RNA samples were collected at each time point from root tissue. Transcript abundance was measured using an ATH1 Arabidopsis genome array and normalized using systematic variation normalization. The raw microarray datasets can be found in the Gene Expression Omnibus under accession numbers GSE84446 and GSE42007. The response to ACC or IAA was reported as Signal Log Ratio (of treatment over the time-matched control for each time point, $log_2$ fold change), and the datasets were filtered to identify transcripts that were above the microarray background, had consistent magnitude and patterns of change and had an SLR>0.5. This resulted in 449 DE genes for the ACC dataset and 1246 DE genes for the IAA dataset. The SLRs of these samples at each time point were used for PaLD community network analyses. For the ACC microarray dataset, we identified 1 duplicate gene ID and used the first instance of this gene in the list (both instances had increased SLRs in response to ACC over time). We also removed the reference gene GAPDH from the list, as this had a non-matching probe ID, leaving us with 447 transcripts to work with (hence the ACC 447, which appears in contradiction to the original publication). Totals of 447 and 1246 transcripts were associated with unique SLRs in the ACC and IAA microarray datasets, respectively, and therefore were used to generate the PaLD networks. However, the expanded list of 1284 genes, which was made by separating multiple gene IDs linked to a single probe on the microarray chip, was used in the matrix table and enrichment analysis for the IAA microarray dataset.

##### Gold Standard Dataset:

The ACC microarray data was reanalyzed in parallel with two additional transcriptomic data sets [62]. Raw data was downloaded and combined for parallel processing using *limma* and other packages in R. Only the 4-hour time point in [50] was used to be consistent between all 3 datasets. Genes were deemed differentially expressed (DE) if they had an absolute fold change greater than 0.5, and an adjusted $p$-value less than 0.05. This meta-analysis identified 139 genes, which we refer to as the *gold standard* of ethylene signaling in roots, which were DE in all three datasets. Because of the parameters defined, only 45 of the gold standard genes were present in the ACC 447. For more detail about the ethylene signaling gold standard genes, see [62].

##### Ethylene RNA-Seq dataset:

The dataset in [51] used a time course to examine ethylene response in Arabidopsis, but with some key differences compared to the dataset in [50]. Plants were treated with ethylene gas (10 μl $l^{-1}$). RNA was then isolated from whole Arabidopsis seedlings grown under different conditions (3 days old, etiolated), rather than isolated root tissue. The time points after treatment were 0, 0.25, 0.5, 1, 4, 12, and 24 hr, which are similar to the time points in [50], but with an additional early time point (0.25 hr), and fewer later time points (additional 2- and 8-

hour time points are in [50]). In addition, RNA-Seq was used to quantify transcripts, rather than microarray.

#### Salt stress RNA-seq dataset:

This time series dataset was generated by growing Col-0 seedlings for 7 d under continuous light and seedlings were transferred to media containing 0 or 100 mM NaCl for 0, 1, 3, 6, 12, 24, 48 and 72 h [53]. RNA was isolated from root samples and RNA-Seq was performed in triplicate. Differential expression analysis was performed using the R package DESeq2 [64] followed by pairwise comparisons to the expression level at 0 h with the threshold of $|\log2(FC)| \geq 1$ and FDR $< 0.1$. The differentially expressed genes were then clustered by the short time-series expression miner (STEM) algorithm using the default settings. We used the top 10 STEM groups in our enrichment analysis. All differentially expressed salt stress genes were used in the PaLD analysis.

#### Supplemental Tables and Figures

*Table S1. Summary of the time series transcriptomic datasets used in this study. For the microarray and RNA-Seq datasets, the processed signals (log ratios) and differentially expressed genes were used in our analysis, respectively.*

| Dataset | Type | Number of DE genes | Treatment | Time course (h) | Clustering method | Accession number |
|---|---|---|---|---|---|---|
| Harkey et al., 2018 | Microarray | 447 | 1 µM ACC | 0, 0.5, 1, 2, 4, 8, 12, 24 | *k*-means | GSE84446 |
| Lewis et al., 2013 | Microarray | 1246 | 1 µM IAA | 0, 0.5, 1, 2, 4, 8, 12, 24 | *k*-means | GSE42007 |
| Chang et al., 2013 | RNA-Seq | 957 | 10 µl $l^{-1}$ ethylene gas | 0, 0.25, 0.5, 1, 4, 12, 24 | DREM | SRA063695 |
| Wu et al., 2021 | RNA-Seq | 3808 | 100 mM NaCl | 0, 1, 3, 6, 12, 24, 48, 72 | STEM | GSE153103 |

*Table S2. Matrix of the overlap between the PaLD and the top 10 k-*means *clusters for the IAA dataset [52]. The sum of the numbers of transcripts in each PaLD or k-means cluster is listed in the last row or column, respectively. The "Other" column indicates transcripts that were not in the largest component or the top ten clusters.*

| | | ***k*-means Clusters** | | | | | | | | | | | |
|---|---|---|---|---|---|---|---|---|---|---|---|---|---|
| | | A | B | C | D | E | F | G | H | I | J | Other | Sum |
| **PaLD Groups** | 1 | 160 | 0 | 0 | 0 | 1 | 0 | 1 | 0 | 0 | 0 | 24 | 194 |
| | 2 | 0 | 113 | 15 | 2 | 0 | 45 | 0 | 6 | 0 | 0 | 65 | 246 |
| | 3 | 3 | 1 | 0 | 83 | 26 | 15 | 0 | 0 | 0 | 0 | 58 | 186 |
| | 4 | 18 | 0 | 8 | 0 | 0 | 0 | 7 | 0 | 0 | 0 | 69 | 102 |
| | 5 | 57 | 0 | 2 | 0 | 1 | 0 | 43 | 1 | 0 | 8 | 81 | 193 |
| | 6 | 32 | 0 | 0 | 7 | 29 | 0 | 1 | 0 | 0 | 1 | 24 | 94 |
| | 7 | 0 | 3 | 0 | 0 | 0 | 0 | 0 | 0 | 18 | 0 | 11 | 32 |
| | 8 | 0 | 12 | 97 | 0 | 0 | 0 | 0 | 36 | 3 | 5 | 79 | 232 |
| | Other | 0 | 1 | 0 | 0 | 1 | 0 | 0 | 0 | 1 | 0 | 2 | 5 |
| | Sum | 278 | 130 | 122 | 92 | 58 | 60 | 52 | 43 | 22 | 14 | 413 | 1284 |

*Table S3. Summary of enrichment results for the 10 largest k-*means *clusters defined in* [52]. *The top three GO annotations with the lowest FDRs were reported (adjusted p-value ≤ 0.05). TFs with targets enriched logFC ≥1 relative to the genome were reported. *This indicates that the group size is larger in the enrichment analysis due to microarray probes recognizing transcripts from more than one gene or smaller due to removal of probes not associated with a gene identifier.*

| ***k*-means cluster** | **Number of transcripts** | **Enriched GO annotations** | **TFs with enriched targets** |
|---|---|---|---|
| **A** | 277* | Megagametogenesis; ribosomal large subunit biogenesis; rRNA processing | None |
| **B** | 130 | Cell division; response to osmotic stress or water deprivation | bZIP28, bZIP51, CAMTA1, ERF014, FAR1, LBD23, LEP, LOB, NLP7, SPL13B, TINY |
| **C** | 122 | Cell division; regulation of cell cycle | None |
| **D** | 91* | Organonitrogen or cellular catabolic process; cell communication | AT5G18450 |
| **E** | 58 | **Auxin-activated signaling pathway**; negative regulation of cellular process; protein ubiquitination | ABF2, AHDP, ANAC042, AT1G47655, BEH3, bHLH31, bHLH46, BMY2, bZIP(2,44,48,66), BZR1, ERF(11,19, 43), ESE3, FAR1, HB9, IDD11, JKD, MGP, MYB98, MYB118, NLP7, PDF2, RAP2.1, RAP2.9 TCP(3, 7, 9, 15, 20) |
| **F** | 60 | Cell wall organization or biogenesis; polysaccharide metabolic process; secondary metabolic process | bZIP22, ERF(3, 34, 38, 43), HB20, RAP2.1, RAP2.9 |
| **G** | 52 | **Auxin-activated signaling pathway**; regulation of hormone levels; response to organic cyclic compound | ANAC079, ASIL2, AT3G25990, AT3G42860, bHLH31, bZIP(2,28,44, 48), FAR1, GT-1, HB20, IDD(1,7,11), MYB(3R4,3R5,63), RAP2.9, WRKY31 |
| **H** | 42* | None | None |
| **I** | 22 | None | ANAC(7,37,38,45,50,53,54,57,58, 70,75,76,78,79,87,92,96,98,101), AT4G26030, MYB(74,80,96,99,121) |
| **J** | 14 | Regulation of biological quality; **response to hormone** or abiotic stimulus | ANAC040, CRC, HB(9,13,24,53), IDD1, IDD7, JKD, MGP, MYB3R1, RVE5, RVE6, SPL13B |

*Table S4. Summary of enrichment results for the ethylene dataset [18]. The top three GO annotations with the lowest FDRs were reported (adjusted p-value ≤ 0.05). TFs with targets enriched logFC ≥1 relative to the genome were reported. We found enriched GO annotations and targets of TFs in 8 and 7 (out of 9) groups, respectively. Moreover, we found 3 groups (groups 4, 6 and 9) with enriched functions related to signaling, with groups 4 and 9 enriched in ethylene signaling.*

| PaLD group | Number of transcripts | Enriched GO annotations | TFs with enriched targets |
|---|---|---|---|
| **1** | 86 | Response to chemical | None |
| **2** | 128 | Carbohydrate or metal ion transport; oxidation-reduction process | None |
| **3** | 106 | Cellular homeostasis; oxidation-reduction process; photoperiodism | ABF2, bZIP(22, 26, 28), LOB, CAMTA1 |
| **4** | 64 | **Ethylene-activated signaling pathway**; negative regulation of signal transduction; tissue development | ABF2, ASIL2, AT1G01250, AT3G11280, AT4G18450, AT4G31060, AT5G18450, bHLH31, bZIP(1, 22, 28, 44, 57), CRF4, DDF1, EIN3, ERF(2, 3, 19 38), ESE1, GAL3, IDD11, JKD, LBD23, LOB13, PUCHI, RAP(2.1, 2.6, 2.9, 2.12), TG, TGA10, TINY, WRKY31 |
| **5** | 106 | Monocarboxylic acid biosynthetic process; oxidation-reduction process; response to nematode | CAMTA5, MYB3R-4, MYB121 |
| **6** | 132 | **Hormone-mediated signaling pathway** | ABF2, AT5G18450, bZIP26, DOF4.2, EIN3 |
| **7** | 148 | Carboxylic acid transport; cell wall polysaccharide biosynthetic process; cellular glucan metabolic process | ESE1, TCP7, TCP15, TINY |
| **8** | 82 | None | ERF10 |
| **9** | 105 | Cellular response to nutrient levels; **ethylene-activated signaling pathway**; protein ubiquitination | bZIP(2, 44, 48), EIN3, GATA-4 |

*Table S5. Matrix of the overlap between the PaLD (horizontal) and the top 10 STEM (vertical) groups* [53] *in the salt dataset. The sum of the transcripts in each PaLD or STEM group is listed in the last row or column, respectively. The "Other" row includes transcripts that were not in the largest component.*

| PaLD Groups | STEM Groups: A | B | C | D | E | F | G | H | I | J | Sum |
|---|---|---|---|---|---|---|---|---|---|---|---|
| *1* | 2 | 0 | 369 | 9 | 131 | 0 | 3 | 3 | 0 | 0 | 517 |
| *2* | 261 | 0 | 12 | 293 | 1 | 0 | 0 | 0 | 0 | 2 | 569 |
| *3* | 0 | 15 | 0 | 0 | 0 | 33 | 0 | 0 | 156 | 101 | 305 |
| *4* | 230 | 2 | 141 | 106 | 17 | 0 | 3 | 0 | 0 | 0 | 499 |
| *5* | 0 | 435 | 0 | 0 | 0 | 202 | 0 | 0 | 4 | 48 | 689 |
| *6* | 2 | 1 | 25 | 2 | 175 | 3 | 198 | 141 | 0 | 0 | 547 |
| *7* | 0 | 258 | 2 | 1 | 0 | 75 | 9 | 32 | 2 | 0 | 379 |
| *8* | 245 | 0 | 0 | 15 | 0 | 0 | 19 | 0 | 0 | 0 | 279 |
| *Other* | 4 | 2 | 4 | 1 | 0 | 1 | 1 | 4 | 4 | 3 | 24 |
| *Sum* | 744 | 713 | 553 | 427 | 324 | 314 | 233 | 180 | 166 | 154 | 3808 |

*Table S6. Summary of enrichment results for the PaLD groups generated from the salt stress RNA-Seq dataset [53]. The top 3 GO annotations with the lowest FDRs were reported (adjusted p-value ≤ 0.05). TFs with targets enriched logFC ≥1 relative to the genome were reported.*

| PaLD groups | Number of transcripts | Enriched GO annotations | TFs with enriched targets |
|---|---|---|---|
| **1** | 517 | Cold acclimation; response to abscisic acid; response to high light intensity | DDF2 |
| **2** | 569 | Auxin-activated signaling pathway; response to red or far red light; transmembrane receptor protein tyrosine kinase signaling pathway | LOB |
| **3** | 305 | Cellular response to hormone stimulus; response to organic cyclic compound; single organism cellular process | AT1G10250, AT4G18450, ERF5, LEP |
| **4** | 499 | Photosynthesis, light harvesting in photosystem I; protein-chromaphore linkage; response to cytokinin | TCP9 |
| **5** | 689 | Maturation of SSU-rRNA from tricistronic rRNA transcript (SSU-rRNA, 5.8s rRNA, LSU-rRNA); ribosomal large subunit biogenesis; ribosomal small subunit biogenesis | None |
| **6** | 547 | Abscisic acid-activated signaling pathway; ethylene-activated signaling pathway; response to chitin | AT1G01250, bHLH46, DDF2, ERF2, GATA-19, LOB, TCP9 |
| **7** | 379 | Defense response to bacterium; protein phosphorylation; response to endoplasmic reticulum stress | None |
| **8** | 279 | None | None |

*Table S7. Summary of enrichment results for the top 10 STEM groups for the salt stress dataset* [53]. *The top 3 GO annotations with the lowest FDRs were reported (adjusted p-value ≤ 0.05). TFs with targets enriched logFC ≥1 relative to the genome were reported. While both PaLD and STEM yielded similar quantities of output for enriched GO annotations, a higher proportion of PaLD groups (5 out of 8, or 63%; Table S6) were enriched in targets of TFs as compared to STEM groups (5 out of 10, or 50%; Table S7). All of the STEM groups that were enriched in targets of TFs had multiple TFs targeting them, while most of the PaLD groups with enriched targets of TFs had one TF targeting them.*

| STEM group | Number of transcripts | Enriched GO annotations | TFs with enriched targets |
|---|---|---|---|
| **A** | 744 | Acyl-coA biosynthetic process; hydrogen peroxide catabolic process; photosynthesis, light harvesting | None |
| **B** | 713 | Cell wall pectin biosynthetic process; response to cadmium ion; tRNA aminoacylation for protein translation | None |
| **C** | 553 | Response to abscisic acid; **response to salt stress**; response to water deprivation | None |
| **D** | 427 | Protein-chromaphore linkage; response to far-red light; response to red light | None |
| **E** | 324 | Abscisic acid-activated signaling pathway; cuticle development; response to desiccation | AT1G01250, ERF2, LBD23, LOB |
| **F** | 314 | Maturation of LSU-rRNA; maturation of SSU-rRNA; protein targeting to mitochondrion | None |
| **G** | 233 | Defense response to bacterium; protein phosphorylation; response to chitin | AT1G01250, AT5G18450, bHLH31, bZIP28, ERF014, LOB, RAP2.1 |
| **H** | 180 | Cell death; defense response; protein autophosphorylation | ANAC054, ANAC079 |
| **I** | 166 | None | AT1G01250, bZIP28, CAMTA1, FAR1, LEP |
| **J** | 154 | Oxoacid metabolic process; response to toxic substance; small molecule biosynthetic process | bZIP22, bZIP26 |

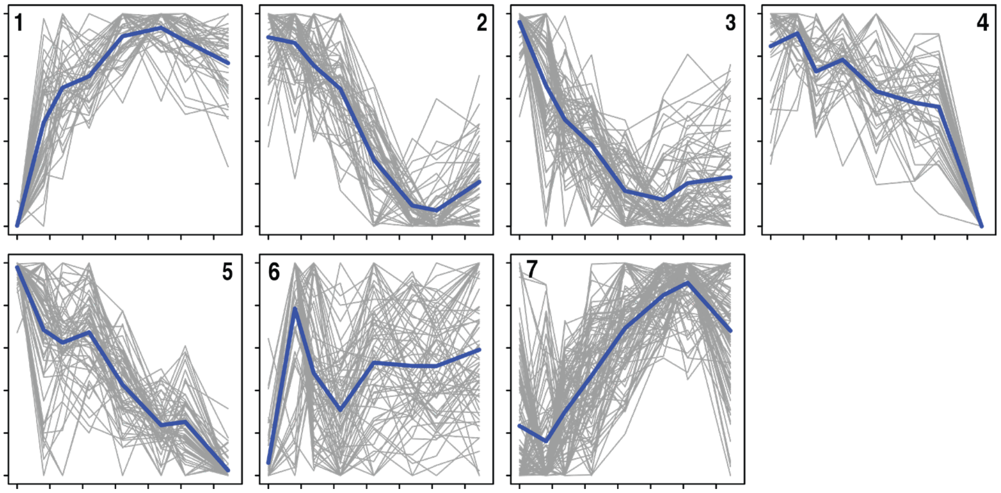


*Fig. S1. The transformed abundance levels before fitting the cubic polynomials for all transcripts in PaLD groups 1–7. Indicated in blue is the average at the respective time points. Note how the curves in each group share fundamental graphical properties concerning relative maxima and minima and intervals of increase/decrease.*

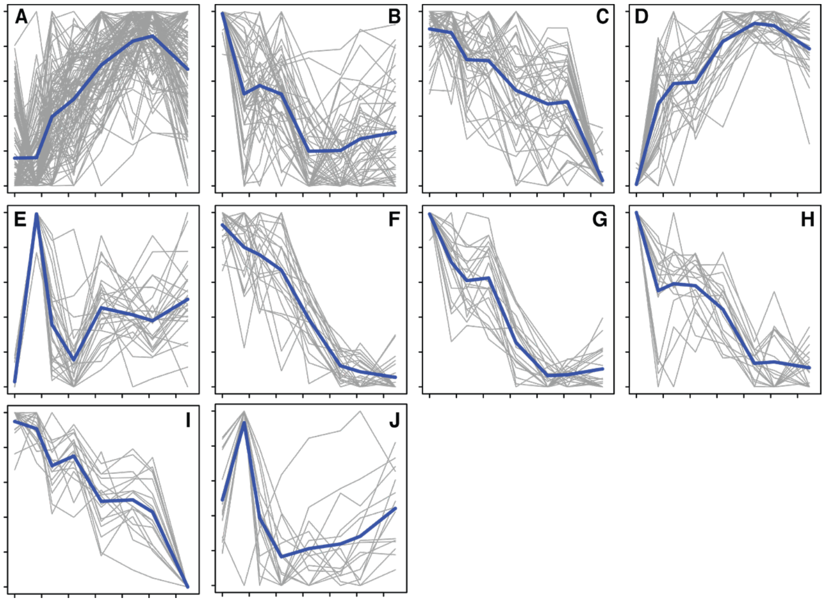


*Fig. S2. The transformed abundance levels before fitting the cubic polynomials for all transcripts in the k-means clusters from Harkey et al. 2018. Indicated in blue is the average at the respective time points. The curve in each cluster share important graphical properties.*

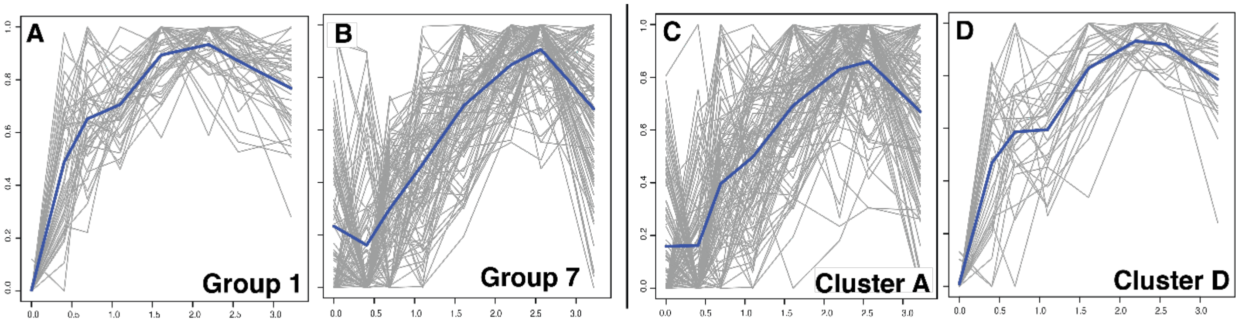


*Fig. S3. provides a plot of the transformed data (prior to fitting of the cubic polynomials) for the transcripts in groups 1 and 7, alongside those for clusters A and D, with similar temporal response patterns. PaLD groups 1 and 7 are over-represented in genes involved in regulation of ethylene signaling and signal transducer activity, respectively. While cluster D is enriched in functional annotation (e.g., regulation of ethylene signaling), the large cluster A, consisting of nearly a third of the transcripts considered in Table 2, has no enrichment in function. This illustrates further the ability of the approach to more effectively group together transcripts based on common biological function. Corresponding plots for the remaining groups and clusters are included in Figs. 9 and 10.*

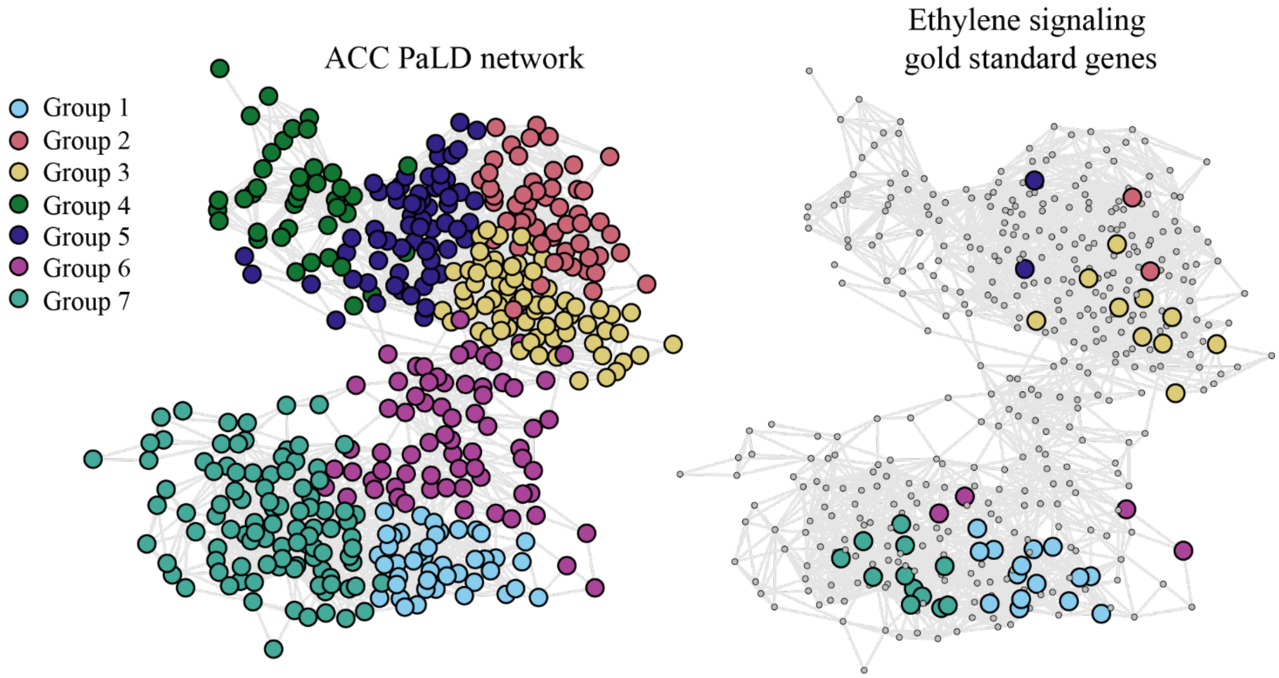


Fig. S4. Overlay of gold standard genes on the PaLD network. The network on the left is the original PaLD network with groups defined by the community detection metric. The network on the right is the PaLD network with the ethylene signaling gold standard genes overlaid on top. The colors correspond to the original PaLD groups (shown on the left). The PaLD algorithm placed the gold standard genes into a small number of groups. Thus, the gold standard genes have similar response curves based on our defined distance function (Eq. (6)). *Of the 447 transcripts used in the PaLD analysis, 45 were part of the group of gold standards. The core ethylene genes were found in 6 of the 7 PaLD groups. Furthermore, areas of the network that did not contain any core ethylene genes could be observed. The distribution of these gold standards across the PaLD groups and k-means clusters are also shown in Tables 2 and 3, which illustrate that they are concentrated in 3 PaLD groups, with 80% found in groups 1, 3, and 7. The partitioning of these gold standard genes into limited PaLD groups is particularly striking, since the temporal response to treatments was not part of the identification of this set of ethylene-responsive transcripts.*

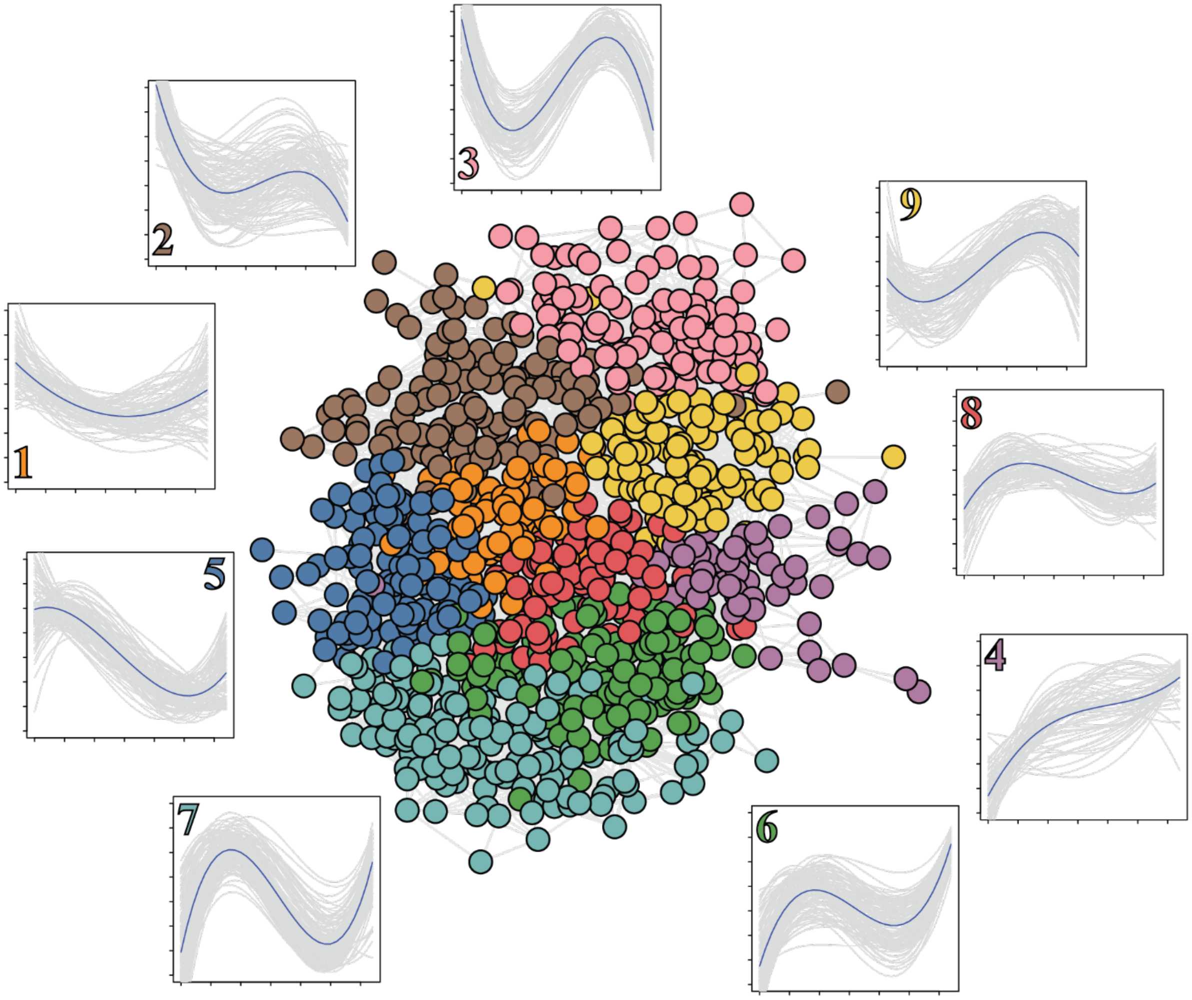


*Fig. S5. PaLD analysis of transcripts with altered gene expression in response to the plant hormone ethylene* [51]*. The network of the 957 transcripts in the ethylene RNA-Seq dataset with each PaLD group having a distinct color is shown in the center. Fiber plots of the PaLD groups illustrating transcript abundance over time color coded to match the network are displayed around structure. Data is considered in log time and has been min-max scaled. Number colors are matched to those in the network. Interestingly, the PaLD network of this dataset did not take on either of the shapes of the PaLD networks of the ACC and IAA microarray datasets. The fiber plots for the PaLD groups conveyed diversity in transcriptional response despite the seemingly high interconnectedness of the groups, and the transcripts in each group are shown in Supplemental Dataset 1.*

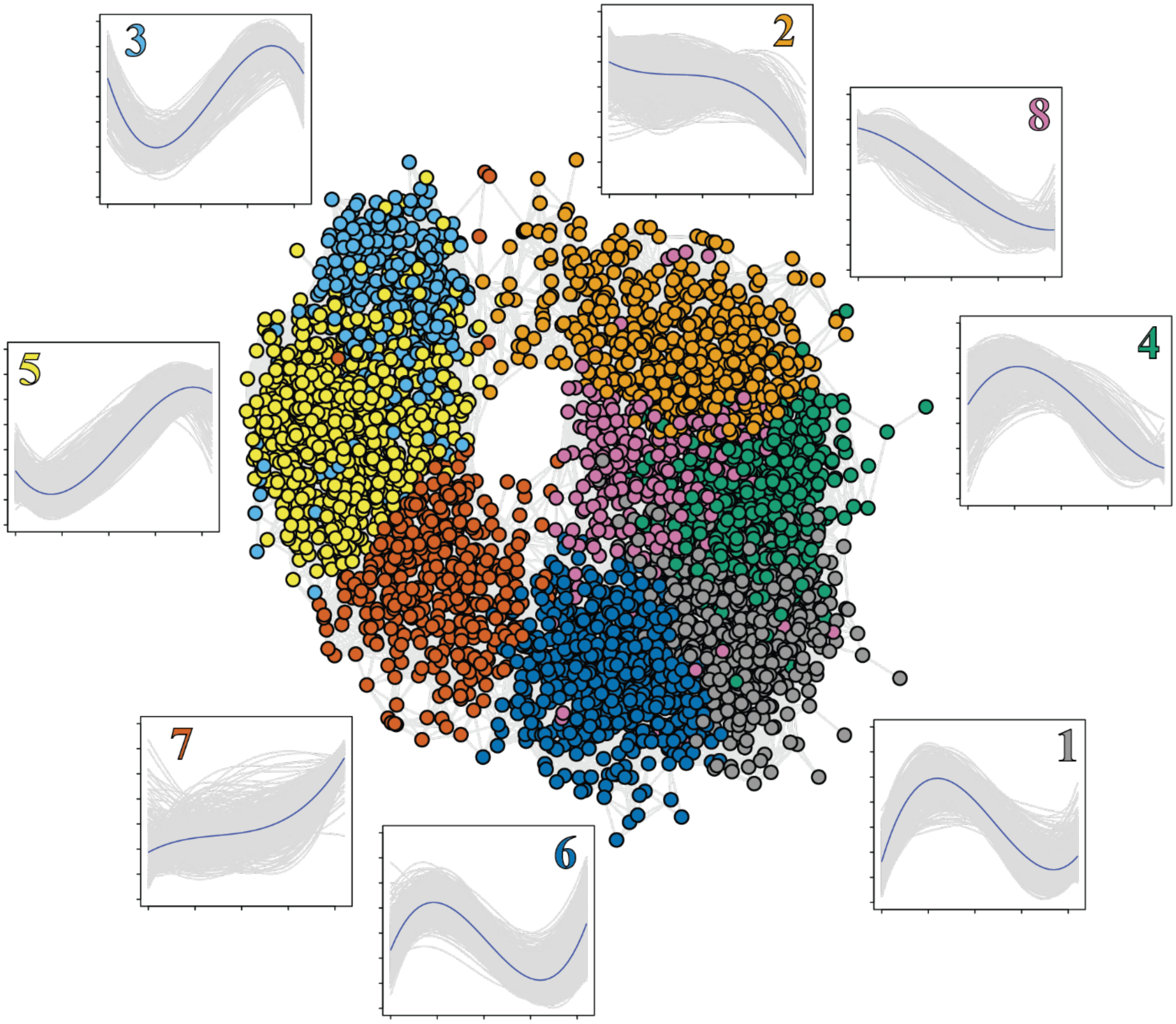


*Fig. S6. PaLD analysis of transcripts with altered gene expression in response to salt stress [53]. These transcripts were derived from an RNA-Seq dataset that had been processed using the short time-series expression miner, or STEM [41,52], which clusters and visualizes short (8 time points or fewer) time series gene expression data. The network of the 3,808 transcripts in the salt stress RNA-Seq dataset is illustrated* with PaLD subgroups having distinct colors. *Fiber plots of the PaLD groups illustrating transcript abundance over time are displayed adjacent to their respective groups in the network. Data is considered in log time and has been min-max scaled. The label for the fiber plots for each group are colored to match the network.*
